# A Single-cell Perturbation Landscape of Colonic Stem Cell Polarisation

**DOI:** 10.1101/2023.02.15.528008

**Authors:** Xiao Qin, Ferran Cardoso Rodriguez, Jahangir Sufi, Petra Vlckova, Jeroen Claus, Christopher J. Tape

## Abstract

Cancer cells are regulated by oncogenic mutations and microenvironmental signals, yet these processes are often studied separately. To functionally map how cell-intrinsic and cell-extrinsic cues co-regulate cell-fate in colorectal cancer (CRC), we performed a systematic single-cell analysis of 1,071 colonic organoid cultures regulated by 1) CRC oncogenic mutations, 2) microenvironmental fibroblasts and macrophages, 3) stromal ligands, and 4) signalling inhibitors. Multiplexed single-cell analysis revealed a stepwise epithelial differentiation landscape dictated by combinations of oncogenes and stromal ligands, spanning from fibroblast-induced Clusterin (CLU)^+^ revival colonic stem cells (revCSC) to oncogene-driven LRIG1^+^ hyper-proliferative CSC (proCSC). The transition from revCSC to proCSC is regulated by decreasing WNT3A and TGF-β-driven YAP signalling and increasing KRAS^G12D^ or stromal EGF/Epiregulin-activated MAPK/PI3K flux. We find APC-loss and KRAS^G12D^ collaboratively limit access to revCSC and disrupt stromal-epithelial communication – trapping epithelia in the proCSC fate. These results reveal that oncogenic mutations dominate homeostatic differentiation by obstructing cell-extrinsic regulation of cell-fate plasticity.

**Highlights:**

- 1,071-condition single-cell transition map of colonic stem cell polarisation regulated by oncogenic and mircoenvironmental cues.
- Fibroblasts polarise WT colonic epithelia towards *Clu*^*+*^ revCSC via TGF-β1 and YAP signalling.
- APC-loss and KRAS^G12D^ drive a *Birc5*^*+*^, *Lrig1*^*+*^, and *Ephb2*^*+*^ proCSC fate via MAPK and PI3K.
- Oncogenic mutations disrupt stromal regulation of epithelial plasticity, trapping cells in the proCSC fate.

## Introduction

The intestinal epithelium comprises multiple cell-types fulfilling the functions of nutrient absorption, waste elimination, and barrier protection [1]. In the healthy colon, a subpopulation of epithelial cells are maintained in a multipotent stem cell state by the pericryptal mesenchymal niche [2]. Stromal fibroblasts secrete paracrine ligands including WNT, EGF, Noggin, and R-Spondin-1 to maintain epithelial stemness and guide differentiation towards secretory and absorptive cells along the crypt [3]. In colorectal cancer (CRC), oncogenic mutations targeting *Apc, Kras, Braf, Smad4*, and/or *Trp53* cell-autonomously induce a crypt-progenitor phenotype in CRC cells [4]. Thus, in both the healthy colon and CRC, a subpopulation of epithelial cells are maintained in a stem-like state – albeit by different mechanisms.

Colonic epithelial stem cells are traditionally described as LGR5^+^ OLFM4^+^ crypt base progenitors [5]. However, recent single-cell studies of intestinal epithelia have identified additional multipotent cell-types, most notably Clusterin (CLU)^+^ ‘revival’ or ‘foetal’ stem cells [6]. Revival stem cells can be induced following tissue damage to repopulate all epithelial cell-types but are otherwise rare in the homeostatic intestine [7]. Revival-like stem cells have also been implicated in CRC initiation [8], can be observed in developed CRC tumours in a patient-specific manner [9], and are emerging as putative drug-tolerant persister cells in CRC [10]. However, how combinations of oncogenic signals and microenvironmental cues regulate the polarisation of epithelia towards traditional or revival stem cells is unclear.

The CRC tumour microenvironment (TME) is a hetero-cellular system where cell-intrinsic oncogenic mutations and cell-extrinsic stromal and immunological signalling cues co-regulate epithelial cancer cells [11]. Stromal ligands and oncogenic mutations can activate common intracellular signalling pathways in colonic epithelia [12].

Canonically, both stromal WNT/R-Spondin-1 ligands and APC-loss hyper-activate β-catenin signalling, whereas EGF and KRAS/BRAF mutations stimulate the MAPK pathway [1]. As a consequence of their overlapping signalling mechanisms, oncogenic mutations must compete with stromal ligands during oncogenesis – yet how cell-intrinsic and cell-extrinsic cues co-regulate epithelial cell-fate remains elusive.

Here we describe a functional single-cell study exploring how cell-extrinsic and cell-intrinsic cues co-regulate colonic epithelial fate. Parallel perturbation analysis of >1,000 heterocellular organoid cultures using single-cell RNA-sequencing (scRNA-seq) and highly-multiplexed thiol-reactive organoid barcoding *in situ* (TOB*is*) mass cytometry (MC) [13] revealed that fibroblasts and oncogenic mutations induce distinct epithelial stem cell-fates in colonic epithelia. We find that fibroblasts polarise epithelia towards slow-cycling CLU^+^ revival stem cells via TGF-β1 and YAP, whereas APC-loss, KRAS^G12D^, and/or exogenous Epiregulin (EREG) shift cells towards a LRIG1^+^ hyper-proliferative fate that is dependent on PI3K signalling. APC-loss and KRAS^G12D^ collaboratively block cell-extrinsic regulation of epithelial plasticity by interrupting stromal-epithelial communication, trapping CRC cells in a cancerous state. Despite the dominance of oncogenes over epithelial plasticity, we find that CRC organoids can still access revival stem cells, but this requires high cell-extrinsic activation of YAP via TGF-β1 in parallel with reduced PI3K signalling.

These results demonstrate that colonic epithelia exist on a continuous differentiation landscape where oncogenic mutations and stromal cues compete for epithelial identity – but oncogenes eventually dominate by blocking the stromal regulation of cell-fate plasticity.

## Results

### Oncogenic and Stromal Cues Differentially Regulate Colonic Epithelia

To directly compare how CRC oncogenic mutations and stromal cells regulate colonic epithelial differentiation, we performed a multivariate scRNA-seq analysis of wild-type (WT), *shApc* (A), *shApc* and *Kras*^*G12D/+*^ (AK), and *shApc, Kras*^*G12D/+*^ and *Trp53*^*R172H/–*^ (AKP) colonic organoids, in monoculture or co-cultured with colonic fibroblasts and/or macrophages (Figure 1A). Fibroblasts are established regulators of intestinal epithelia [14] and macrophages are the most profuse leukocytes in the colon [15]. WT epithelia cultured with exogenous WNT3A, EGF, Noggin, and R-Spondin-1 (WENR) (commonly used to grow colonic organoids) were included as a defined mesenchymal niche factor control.

**Figure 1.**
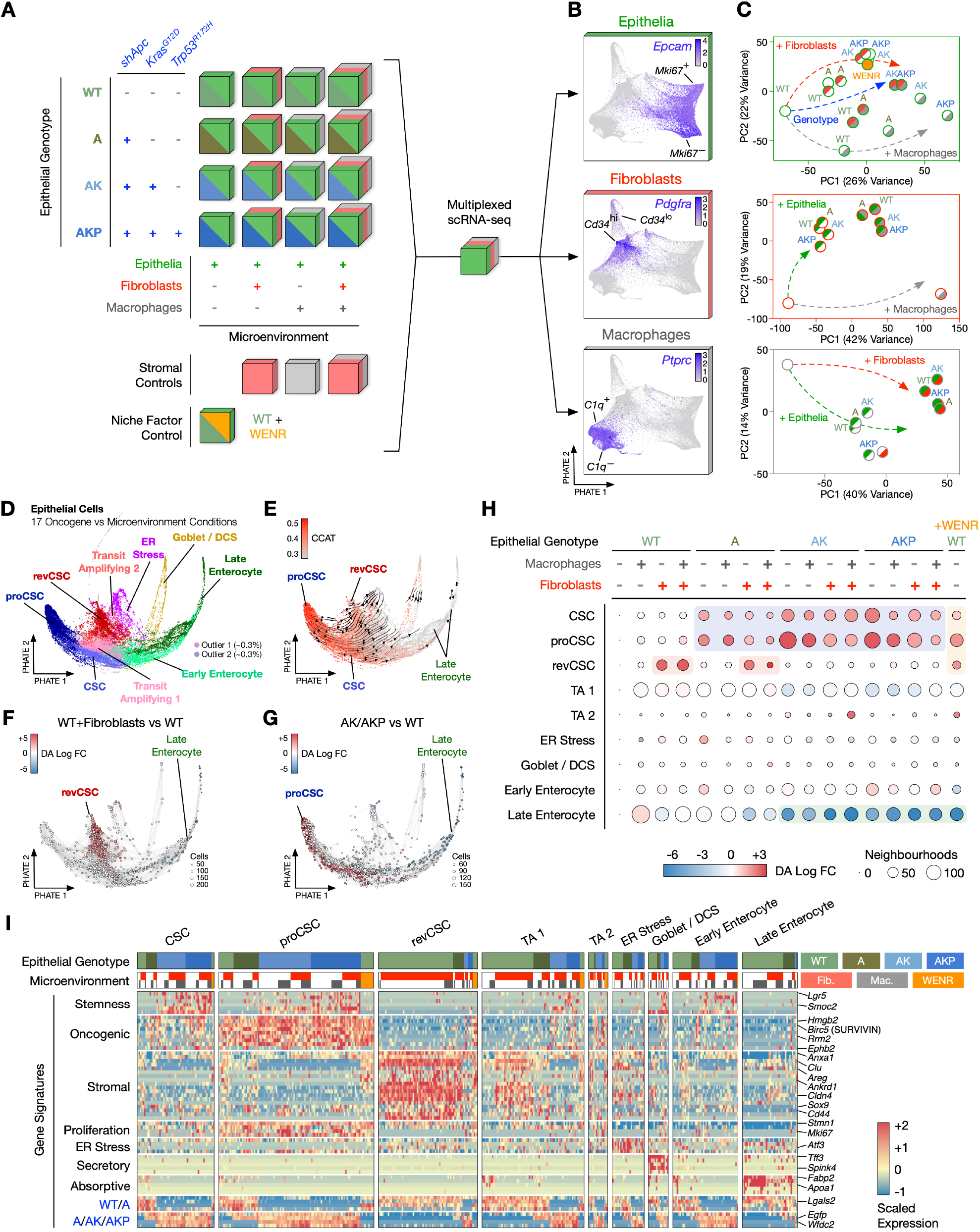
Oncogenes and Fibroblasts Differentially Regulate Colonic Epithelia. **A)** Multivariate scRNA-seq experimental design. WENR ligands were removed from all experimental conditions except for the niche factor control to ensure cell-cell signalling was not dominated by exogenous recombinant proteins (see Methods). **B)** Single-cell PHATE embedding illustrating epithelial cells, fibroblasts, and macrophages. **C)** PCAs of epithelial, fibroblast, and macrophage transcriptomes regulated by by organoid genotype and microenvironment. **D)** PHATE embedding of 29,452 epithelial cells from the 17 organoid conditions coloured by cell-type clusters. **E)** Epithelial PHATE coloured by CCAT score and overlaid with velocity streams (arrows). **F)** Epithelial PHATE overlaid with differentially abundant (DA) neighbourhoods in WT organoid + fibroblast co-cultures compared with WT organoid monocultures. **G)** Epithelial PHATE overlaid with DA neighbourhoods in AK/AKP organoid monocultures compared with WT organoid monocultures. **H)** Dot plot of epithelial clusters across organoid cultures coloured by log fold-change (Log FC) in neighbourhood abundance and sized by the number of neighbourhoods detected. **I)** Gene expression signatures of epithelial clusters. WENR, WNT3A, EGF, Noggin, and R-Spondin-1. CSC, colonic stem cell. proCSC, hyper-proliferative CSC. revCSC, revival CSC. DCS, deep crypt secretory cell. CCAT, correlation of connectome and transcriptome. TA, Transit amplifying cell.

Following scRNA-seq, epithelial cells, fibroblasts, and macrophages were resolved by Leiden clustering [16], visualised by PHATE (Potential of Heat-diffusion for Affinity-based Trajectory Embedding) [17] (Figure 1B), and cell-type-specific transcriptional changes were summarised by principal component analysis (PCA) (Figure 1C). Epithelial transcriptomes are differentially regulated by both CRC mutations (PC1, 26%) and microenvironmental cues (PC2, 22%), with A, AK, and AKP mutations progressively dysregulating their transcriptomic profiles. However, we found fibroblasts can only regulate WT and A epithelial cells (Figure 1C). Although WENR ligands are thought to mimic a healthy stromal niche [18], WT organoids + WENR ligands transcriptionally align with AK mutant organoids (not WT+fibroblasts as might be expected), indicating this widely used colonic organoid culture media induces a partial CRC-like transcriptome in WT epithelia (Figure 1C).

Colonic fibroblasts clustered into CD34^hi^ and CD34^lo^ subpopulations mimicking *in vivo* stromal heterogeneity [19, 20] (Figure S1A). CD34^hi^ and CD34^lo^ fibroblasts did not differentially regulate colonic epithelia (Figure S1B) and were subsequently treated as a heterogenous mesenchymal population. We found fibroblast and macrophage transcriptomes were only regulated by co-culture with heterotypic cells but not altered by epithelial genotypes (Figures 1C, S1C-D).

### Oncogenic Mutations and Fibroblasts Polarise Epithelia Towards Distinct Stem Cell-Fates

Epithelial cells from all conditions were integrated by reciprocal PCA (RPCA) [16], projected onto a shared PHATE embedding, and clustered into multiple cell-fates, including stem populations, transit amplifying (TA) cells, cells under ER stress, goblet and deep crypt secretory (DCS) cells, and early or late enterocytes (Figure 1D). Stem clusters contain high signalling entropy (indicative of pluripotency) [21] and act as origins for RNA velocity streams [22] that transition towards differentiated cells (Figures 1E, S2E).

Differential abundance testing [23] of co-culture and CRC monoculture conditions against WT monocultures revealed that fibroblasts, macrophages, and CRC mutations have markedly different effects on epithelial cell-fate determination (Figure 1F-H). Fibroblasts enrich a distinct stem cell population characterised by high expression of epithelial progenitor genes *Clu, Sox9, Cd44*, and *Cldn4* (Figures 1I). These fibroblast-induced stem cells are transcriptionally similar to ‘foetal’ [24, 25] or ‘revival’ stem cells (revSCs) [7] of the small intestine (S2A) and are hereafter referred to as ‘revival colonic stem cells’ (revCSC).

In contrast, A, AK, and AKP mutations progressively polarise epithelia towards a hyper-proliferative colonic stem cell-fate, hereafter named proCSC (Figure 1G, H). proCSCs express *EphB2, Birc5* (*Survivin*), *Lrig1, Hmgb2*, and *Rrm2* and are highly mitotic (*Stmn1*^*+*^, *Mki67*^*+*^, and *Ccnb1*^*+*^) (Figure 1I). In addition, proCSCs are transcriptionally comparable to stem cells observed in mouse and human CRC (Figure S2A). Both revCSC and proCSC are present in WT organoids at low levels alongside traditional *Lgr5*^*+*^ colonic stem cells, hereafter named CSC (Figure S2B). We found CSC are also enriched by A, AK, and AKP genotypes, but to a lesser extent than proCSC, and CSC gene signatures are less common in CRC (Figure S2A).

We found that fibroblasts can only induce revCSC in WT and *shApc* epithelia, but not when cells contain both *shApc* and *Kras*^*G12D/+*^ (Figure 1H). Conversely, proCSCs are enriched in all A, AK, and AKP organoids irrespective of fibroblasts or macrophages, suggesting oncogenic mutations are dominant over microenvironmental signalling. WENR ligands hyper-polarise WT epithelia towards all stem and TA cell-types, with very few cells retaining secretory or absorptive identities (Figures 1H-I, S2B). WT epithelia also show higher RNA velocity vector lengths relative to CRC cells (Figure S2C-D), suggesting that oncogenic mutations reduce epithelial plasticity. While macrophages can alter epithelial gene expression (Figure 1C), macrophages do not regulate the abundance of epithelial cell-types (Figure 1H). In summary, multivariate scRNA-seq revealed that fibroblasts, CRC mutations, and WENR ligands polarise epithelia towards a de-differentiated progenitor state – with fibroblasts and oncogenes inducing distinct revCSC and proCSC fates.

### WNT3A Polarises Epithelia to revCSC and Oncogenic Mutations to proCSC

Multivariate scRNA-seq demonstrated that cell-extrinsic ligands and cell-intrinsic mutations differentially regulate epithelial cell-fate, but could not describe how individual ligands and mutations co-regulate differentiation. To functionally explore epithelial polarisation, we performed a highly-multiplexed TOB*is* MC [12] combinatorial study focusing on the three axes hypothesised to regulate epithelial cell-fate: 1) microenvironment (+/- fibroblasts), 2) stroma-mimicking ligands (+/- WNT3A, +/- EGF, +/- Noggin, +/- R-Spondin-1), and 3) oncogenic mutations (+/- *shApc*, +/- *Kras*^*G12D/+*^, +/- *Trp53*^*R172H/–*^) (Figure 2A). Each organoid culture was performed in triplicate, barcoded *in situ* using 126-plex TOB*is* [13], pooled, dissociated into single cells, stained with a panel of 45 rare-earth metal-labelled antibodies (spanning epithelial differentiation markers identified by scRNA-seq, cell-state markers, and PTM signalling nodes [12]) (Table S1), and analysed by MC. Following debarcoding [26], QC, and cell-type-specific gating, we obtained 6 million cells from 390 organoid/fibroblast cultures (570 cell-type-specific single-cell datasets) (Figure 2B-D).

**Figure 2.**
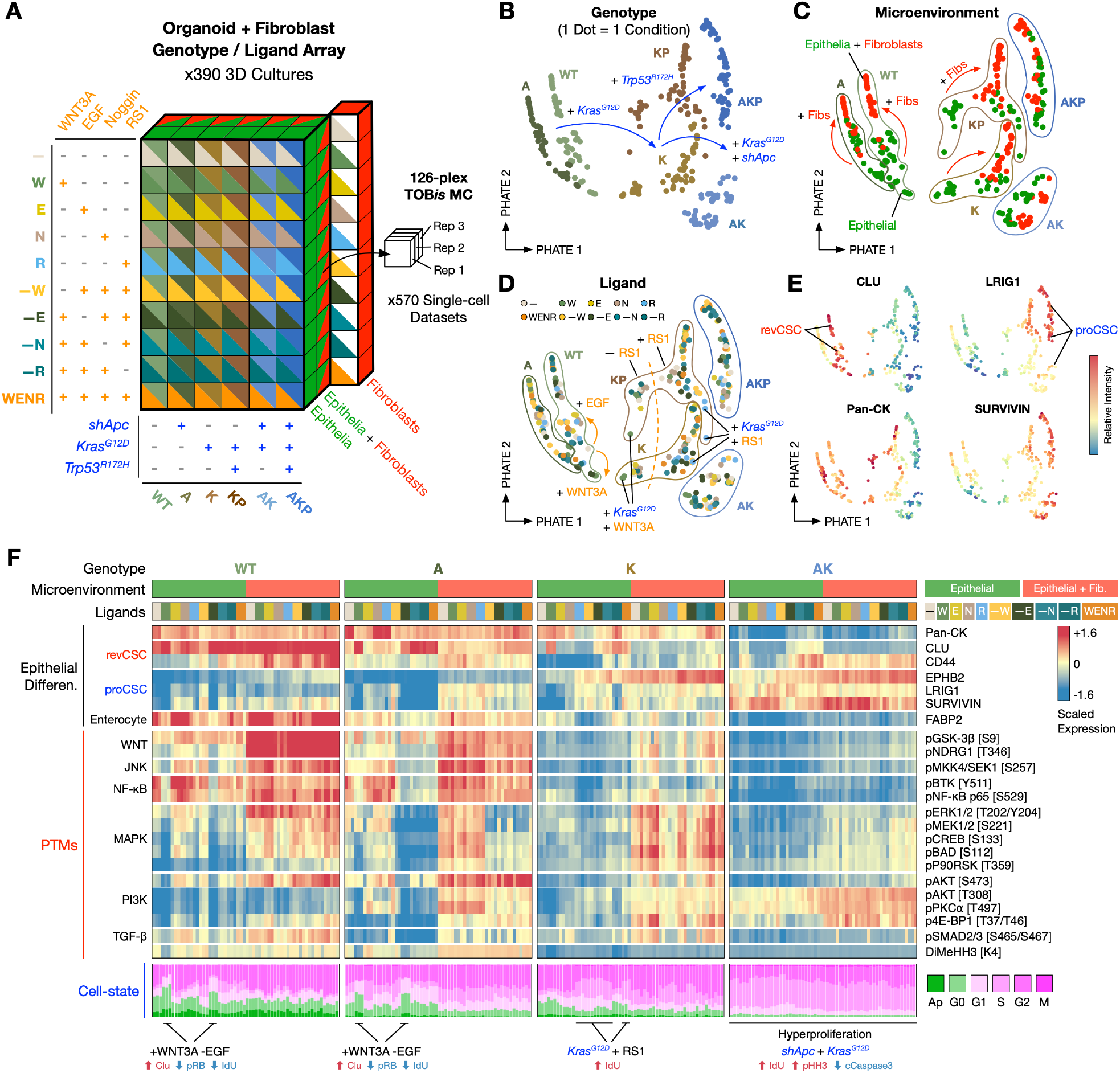
Regulation of Colonic Epithelia by Epithelial Genotypes, Fibroblasts, and WENR Ligands. **A)** TOB*is* MC multidimensional array comprising epithelial genotypes, fibroblasts, and WENR ligands (570 single-cell datasets). **B-E)** EMD-PHATE of 360 organoid cultures coloured by genotype, microenvironment, WENR ligands, and EMD scores of epithelial cell markers. One dot = one condition. **F)** Relative expression of epithelial markers, PTMs, and cell-state markers regulated by genotypes, fibroblasts, and/or WENR ligands. One column = one condition. MC, mass cytometry. Fib., fibroblasts. Ap., Apoptotic.

In agreement with scRNA-seq, analysis of 360 epithelial single-cell profiles confirmed that fibroblasts induce CLU^+^ revCSC in WT epithelia (Figures 2C,E, S3A), whereas oncogenic mutations induce hyper-proliferative LRIG1^+^, EPHB2^+^, and SURVIVIN^+^ proCSC while blocking access to revCSC (Figures 2B, E, S3A).

The effect of WENR ligands on epithelial differentiation is highly dependent on genotype (Figure 2D, F). For example, when WT or A organoids are treated with R-Spondin-1 alone, no distinct shift in cell-signalling or cell-state is observed. However, when K (*Kras*^*G12D/+*^) or KP (*Kras*^*G12D/+*^, *Trp53*^*R172H/–*^) cells are treated with R-Spondin-1, they undergo a dramatic S-phase entry and phenocopy AK and AKP genotypes (Figures 2F, S3B).

This suggests that KRAS^G12D^ fundamentally rewires how epithelial cells respond to canonical β-catenin signalling (via stromal R-Spondin-1 or APC-loss) to bias epithelia towards proCSC. By contrast, WNT3A upregulates CLU in WT, A, K, and KP epithelia but only shows a very minor effect on cells containing both *shApc* and *Kras*^*G12D/+*^ (Figures 2F, S3C), indicating that APC-loss and oncogenic KRAS dominate over epithelial response to exogenous cues, blocking access to revCSC and entrapping epithelia in the proCSC fate.

Despite their origin as stroma-mimicking cues, we found that WENR ligands regulate epithelia very differently from fibroblasts (Figure S3D-F). Purified WNT3A enriches quiescent revCSCs with low mitogenic PTM signalling activity. Conversely, fibroblasts induce SOX9^+^, pRB [S807/S811]^+^ revCSCs with high levels of MAPK (pERK1/2 [T202/Y204], pMKK3/6 [S189/S207], pMAPKAPK2 [T334], and pP90RSK [T359]) and TGF-β (pSMAD2/3 [S465/S467]) signalling (Figure S3F). This suggests that fibroblast-induced revCSCs are distinct from those regulated by WNT3A alone and that the communication between stromal and epithelial cells is more diverse than just WENR ligands.

### Oncogenic Mutations and Stromal Ligands Regulate Epithelia Across a Continuous Differentiation Trajectory

To understand how organoid monocultures are regulated by WENR ligands, we analysed WT, A, K, AK, KP, and AKP organoids treated +/- WNT3A, EGF, Noggin, and R-Spondin-1 (180 single-cell profiles). This analysis revealed that colonic epithelial differentiation exists on a multivariate continuum where stromal ligands and oncogenic cues compete for epithelial fate (Figure 3A). We observed a clear fate-transition trajectory of epithelial differentiation dictated by oncogenes and ligands, spanning from WNT3A-driven WT revCSC, through an equilibrium of balanced stem cell identities and enterocyte differentiation, to oncogene-dominant proCSC (Figures 3A, S4A-B). Crucially, WNT3A can drive epithelia towards the revCSC fate when only one oncogenic-driver is present, but the combination of APC-loss and KRAS^G12D^ traps epithelia in the proCSC state that is largely unresponsive to all WENR ligands (Figure 3A).

**Figure 3.**
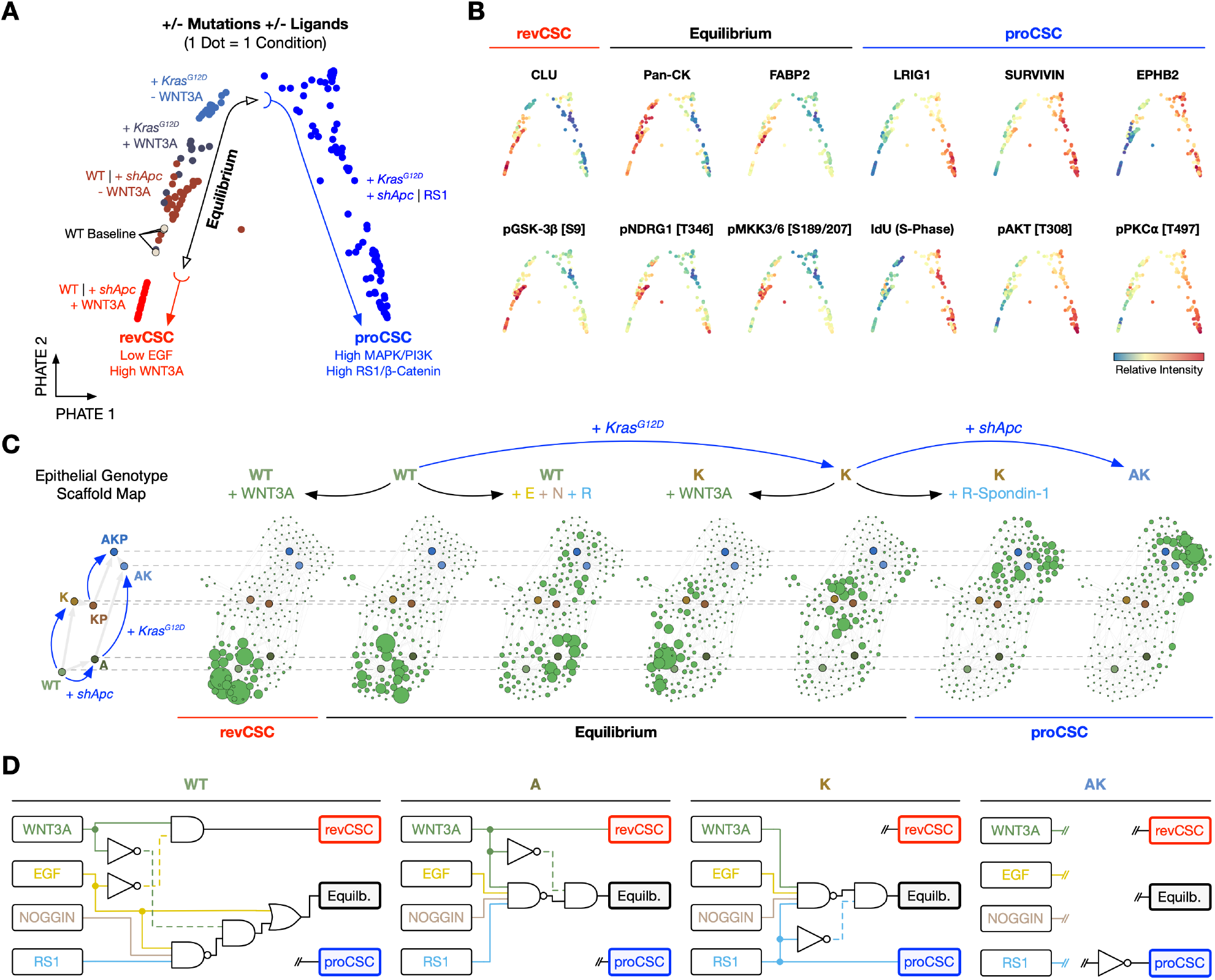
Stepwise Transition from revCSC to proCSC Regulated by Oncogenic Mutations and Ligands. **A)** EMD-PHATE of 180 organoid single-cell datasets regulated by epithelial genotype and WENR ligands. One dot = one condition. **B)** EMD-PHATE coloured by EMD scores of revCSC, equilibrium, and proCSC markers. **C)** Epithelial genotype scaffold maps with organoid monoculture landmarks and genotype+ligand overlays. **D)** Boolean logic models for genotype-specific regulation of colonic stem cells (CSC) by WENR ligands. revCSC, revival CSC. Equilb., equilibrium. proCSC, hyper-proliferative CSC. RS1, R-Spondin-1.

We found that the regulation of revCSC by WNT3A is also heavily influenced by parallel EGF signalling. For example, WNT3A alone leads to quiescent CLU^+^ revCSC in WT epithelia, but if WNT3A is combined with EGF, cells maintain cell-cycle activity and achieve an equilibrium of stem identities (Figures 2F, S4B). The transition between revCSC and equilibrium can be clearly observed across a WNT3A vs EGF gradient and fine-tuned by altering the ratio between EGF and WNT3A concentrations (Figure S4C-F). This suggests that the access to revCSC is controlled by competing signalling flux downstream of WNT3A and EGF.

Consistent with the hypothesis that revCSC and proCSC are regulated by different signalling pathways, TOB*is* MC demonstrated that revCSCs have low cell-cycle activity and high pGSK-3β [S9], whereas epithelia in the equilibrium state display activated pNDRG1 [T346] and pMKK3/6 [S189/S207]. In contrast, proCSC lose cytokeratin expression and have very high levels of PI3K signalling (e.g. pAKT [T308], pPKCα [T497], and p4E-BP1 [T37/T46]) (Figure 3B). The continuous regulation of epithelia by CRC mutations and ligands can be orthogonally depicted in a genotype-anchored scaffold map [27], where revCSC-enriched WT+WNT3A transition into proCSC-dominant K+R-Spondin-1 and AK conditions in a stepwise manner (Figure 3C). The regulation of epithelial stem cell-fate by WENR ligands can be described by simple genotype-specific Boolean logic models (Figure 3D). These models reveal that while WT epithelia are highly sensitive to cell-extrinsic reprogramming, *shApc* and *Kras*^*G12D/+*^ progressively limit epithelial plasticity and cell-intrinsically trap epithelia in the proCSC fate.

### Oncogenic Mutations Inhibit Fibroblast-Epithelia Signalling

As epithelial differentiation cannot be regulated by fibroblasts in the context of *shApc* and *Kras*^*G12D/+*^ (Figures 1H, 2C), we hypothesised oncogenic mutations might disrupt stromal-epithelial signalling. To test this, we performed ligand-receptor cell-cell communication analysis [28] of WT, A, AK, and AKP organoid+fibroblast co-culture scRNA-seq datasets.

Given their established role in microenvironmental cell-cell communication, fibroblasts unsurprisingly demonstrate high ‘outgoing’ signalling (i.e., express numerous ligands and extracellular matrix (ECM) components). By contrast, WT epithelia display a dominant ‘incoming’ signalling potential (i.e., express many receptors) (Figure 4A). This dichotomy suggests that heterocellular signalling in the healthy colon is largely unidirectional from fibroblasts to epithelial cells. We found that revCSC and the closely affiliated TA 1 and TA 2 clusters are responsible for much of the ‘incoming’ signalling potential of WT epithelia, indicating these cell-types are hypersensitive to cell-extrinsic regulation by fibroblasts. In contrast, proCSC are the least receptive of all epithelial cells, suggesting proCSC are more reliant on cell-intrinsic signalling (Figure 4A).

**Figure 4.**
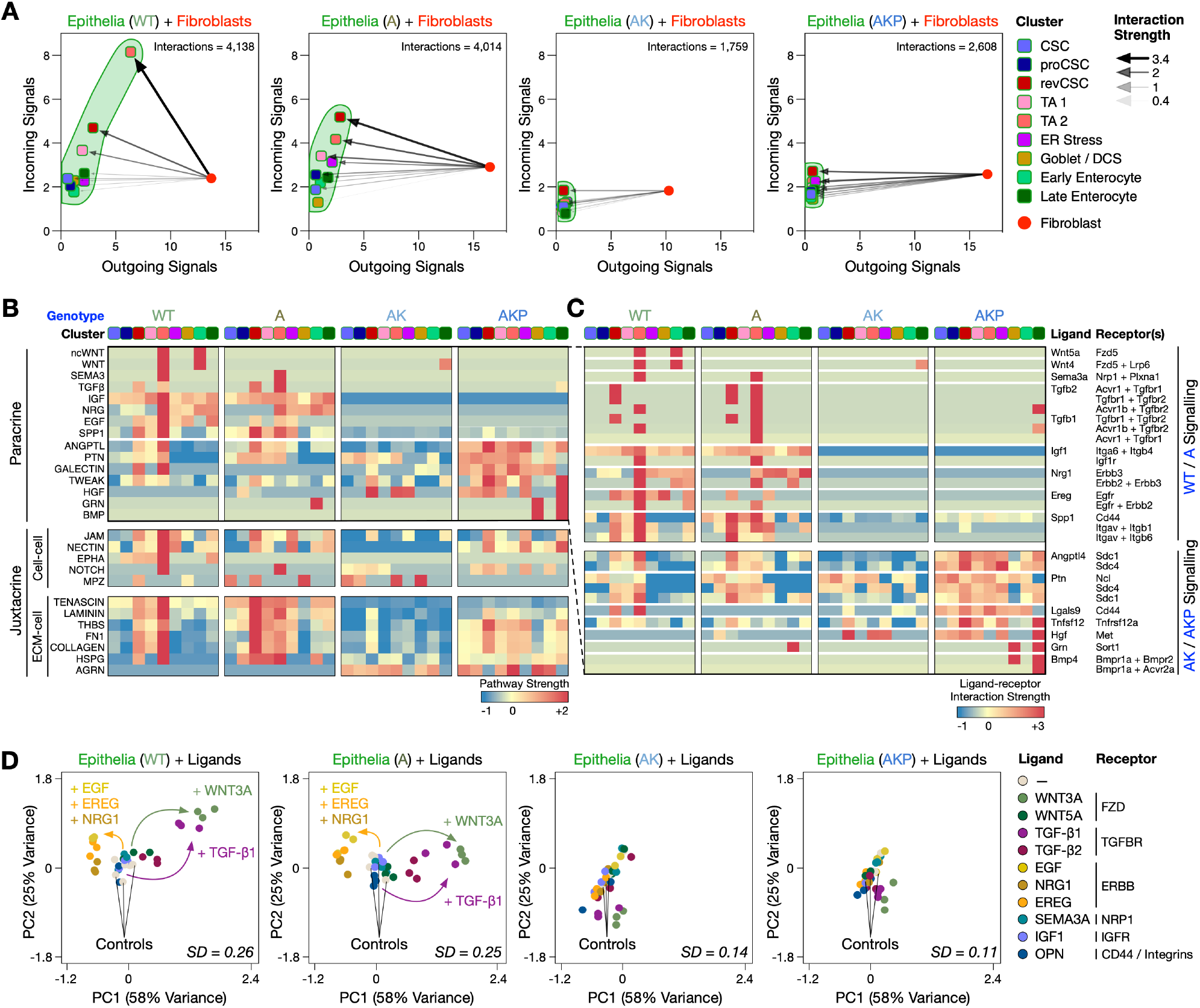
Oncogenic Mutations Disrupt Stromal-epithelial Communication. **A)** Outgoing and incoming communication probability (interaction strength) from fibroblasts to epithelia across organoid genotypes. **B-C)** Predicted paracrine and juxtacrine communication summarised at the pathway and ligand-receptor interaction level. **D)** EMD-PCA of epithelial regulation by exogenous ligands across the genotypes (138 single-cell datasets). One dot = one condition. SD, standard deviation of the distribution of EMD scores for each genotype.

Cell-cell communication analysis revealed that fibroblasts form putative paracrine and juxtacrine interactions with WT and A cells, which are often lost in AK and AKP genotypes (Figure 4B). For example, WT and A organoids show intact NRG1, EREG, IGF, and TGF-β signalling with fibroblasts, but these cell-cell interactions are undetectable in AK and AKP cells, due to the down-regulation of epithelial signal receptors (Figures 4B-C, S5A-C).

Ligand-receptor analysis is increasingly used to generate putative cell-cell communication models in heterocellular systems [29], yet these computational hypotheses are rarely experimentally validated. To functionally test how oncogenic mutations regulate stromal-epithelial communication, we performed a systematic TOB*is* MC study of epithelial differentiation, cell-state, and PTM signalling in WT, A, K, KP, AK, and AKP organoids treated with stromal ligands identified by ligand-receptor analysis as WT homeostatic regulators (WNT5A, SEMA3A, TGF-β1, TGF-β2, IGF, NRG1, EREG, and OPN (*Spp1*)) (Figure 4B-C).

Single-cell MC analysis of 204 organoid cultures revealed that WT and A epithelia can be polarised towards revCSC by WNT3A or TGF-β1, whereas ERBB signalling via EGF, EREG, or NRG1 pushed cells towards the proCSC fate. This suggests that stem cell polarisation can be recapitulated by fibroblast-secreted ligands independent of stromal-epithelial contact or fibroblast-driven ECM remodelling. In contrast, ligands fail to regulate epithelia containing both *shApc* and *Kras*^*G12D/+*^ (Figures 4D, S5D-F). The resistance to external signalling cues of AK/AKP epithelia mimics the diminishing stromal-epithelial communication predicted by ligand-receptor analysis (Figure 4A-C) and is reminiscent of their unresponsiveness to WENR ligands (Figure 3D). Collectively, this analysis suggests that the combination of APC-loss and oncogenic KRAS^G12D^ decouple epithelial cells from homeostatic intercellular signalling – with CRC cells becoming ‘bad listeners’ in the tissue microenvironment.

### revCSC and proCSC are Regulated by Competing Signalling Pathways

As epithelial cells are co-regulated by cell-intrinsic and cell-extrinsic cues across an integrated differentiation trajectory, we hypothesised different signalling pathways might compete to control epithelial cell-fate. To determine the signalling hubs regulating revCSC and proCSC polarisation, we performed an extensive single-cell cuesignal-response perturbation assay spanning: 1) CRC oncogenic mutations (*shApc* and *Kras*^*G12D/+*^), 2) stem cell polarisation ligands (WNT3A, EREG, and TGF-β1), and 3) inhibitors targeting: β-catenin (ICG-001), GSK-3β (CHIR99021), MEK (Trametinib), PI3K (GDC-0941), FAK (PF-573228), SRC (Dasatinib), YAP (CA3), and SMAD3 (SIS3) (Figure 5A).

**Figure 5.**
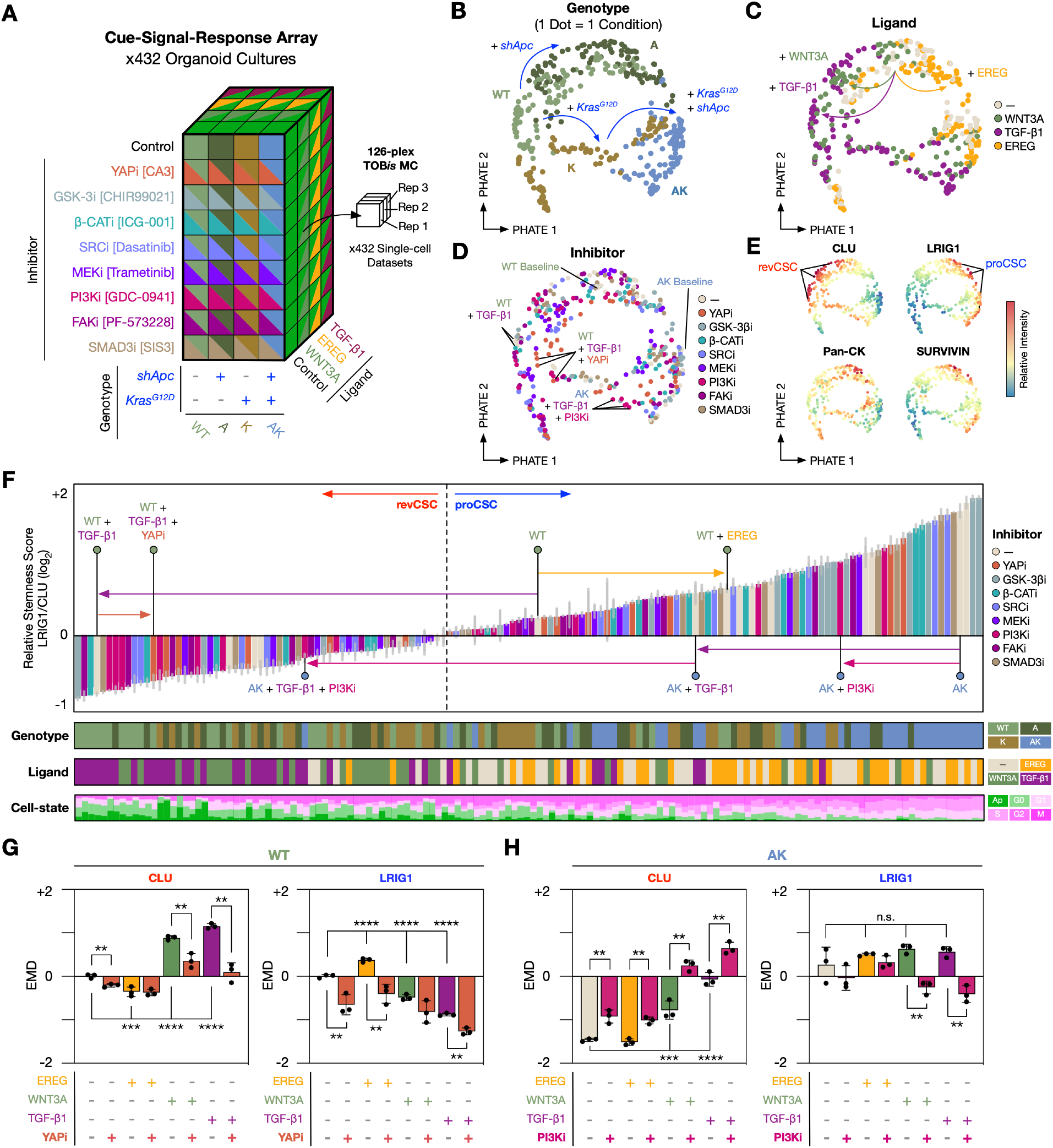
revCSC and proCSC are Regulated by Competing Signalling Nodes. **A)** Cue-signal-response organoid array experimental design. **B-E)** EMD-PHATE of 432 organoid cultures coloured by genotype, ligand, inhibitor, and EMD scores of epithelial cell markers. One dot = one condition. **F)** Ranked relative stemness score (log_2_ -transformed single-cell expression ratio between LRIG1 and CLU) across all conditions in the cue-signal-response array annotated by epithelial genotypes, exogenous ligands, and organoid cell-state. Error bars = SD. **G)** EMD scores for CLU and LRIG1 across WT organoid culture conditions. **H)** EMD scores for CLU and LRIG1 across AK organoid culture conditions. (**, *p* < 0.01; ***, *p* < 0.001; ****, *p* < 0.0001. n.s., not significant. Ordinary one-way ANOVA with Holm-Šídák’s multiple comparisons test between untreated and ligand controls. Two-tailed unpaired *t* -test for inhibitor treatments). Error bars represent SD.

Analysis of 432 single-cell MC organoid profiles confirmed that WT, A, and K epithelia can be polarised towards revCSC by WNT3A or TGF-β1 and to proCSC by EREG. However, organoids containing both *shApc* and *Kras*^*G12D/+*^ showed limited response to ligands and largely retained their proCSC identify (Figures 5B-E, S6A-B). While the ligand-effect is genotype-specific, signalling inhibitors can disrupt the polarisation of proCSC and revCSC across all genotypes, with several interesting examples of ligands and inhibitors collaborating to regulate epithelial cell-fates (Figures 5D, S6D-K).

To rank the polarisation of proCSC and revCSC by genotypes, ligands, and inhibitors across a shared regulation landscape, we established a relative stemness (RS) score by calculating the single-cell expression ratio between LRIG1 and CLU for each organoid culture (Figure 5F). In this space, WT+TGF-β1 have a low RS score, indicating enrichment of CLU^+^ revCSCs, whereas AK have a high RS score and are dominated by LRIG1^+^ proCSCs (Figures 5F, S6C). The differential polarization of proCSC and revCSC can therefore be captured using shifts in the RS score (Figure 5F).

Surprisingly, SMAD inhibition did not alter TGF-β1 regulation of revCSC (Figure S6F). However, both TGF-β1 and WNT3A regulation of revCSC could be partially reversed by YAP inhibition, suggesting revCSC is a YAP-dependent cell-fate (Figures S2A, 5F-G, and S6G). In contrast, although treating AK organoids with either TGF-β1 or PI3Ki alone caused a decrease in RS score (with the epithelial population still dominated by proCSCs), we found treatment of AK organoids with PI3Ki and TGF-β1 enabled epithelia to enter a revCSC-dominant state.

This suggests that CRC cells can access revCSC, but the transition requires high TGF-β1 and low PI3K signalling (Figures 5F, 5H, and S6I). Collectively, the single-cell cue-signal-response perturbation array revealed that colonic stem cell plasticity is generally resilient, but cells can transition between revCSC and proCSC by re-balancing competing signalling flux in YAP, PI3K and MAPK pathways, even in CRC organoids. We found that YAP is a central regulator of revCSC, while PI3K and MAPK are important for maintaining the proCSC identity.

### Single-cell Landscape of Colonic Epithelial Cell-fate Plasticity

In 1957, C.H. Waddington published his famous illustration of cellular differentiation, depicting pluripotent cells rolling down a landscape into valleys of terminal differentiation [30]. While an evocative metaphor in developmental biology, this conceptual model has not been clearly demonstrated with real data. However, recent computational advances in global-structure embeddings [17], differentiation potency metrics [21], and local differentiation-rate predictions [22] now provide the component elements to reconstruct Waddington-like embeddings from scRNA-seq data.

To visualise single-cell colonic epithelial differentiation on a Waddington-like landscape, we combined the global cellular relationships captured by PHATE [17] as ‘longitude and latitude’ axes, with an integrated Valley-Ridge (VR) score to represent pluripotent ‘altitude’. The VR score is defined as the sum of two components per cluster: CCAT signalling-entropy [21] and RNA velocity [22]. At a cluster’s centre, the VR score is solely determined by the median CCAT. However, the VR scores at the cluster periphery were augmented by weighting the inverse of RNA velocity component and the scaled distance from the cluster centre to model rates of local transcriptional change. This method reconstructs a data-driven estimate of Waddington-like landscapes where the altitude captures the differentiation potential of a cell population, with the valley-ridge topology delineating local plasticity (Figure 6A).

**Figure 6.**
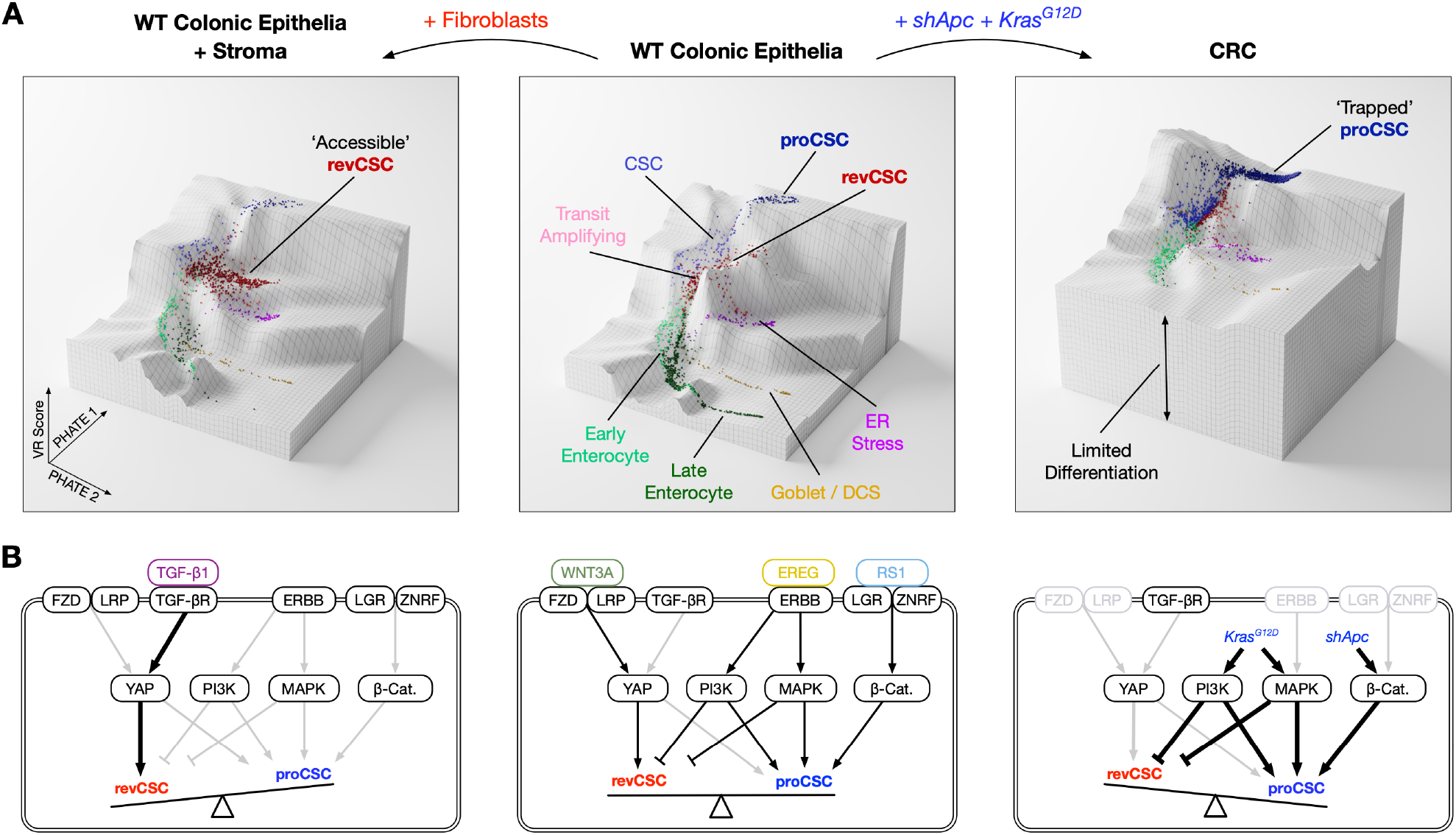
Fibroblast- and Oncogene-driven Waddington-like Single-cell Landscapes. **A)** Integrating PHATE and Valley-Ridge (VR) score enables Waddington-like embeddings of scRNA-seq data. Landscapes illustrate how WT epithelia differentiate from high signalling-entropy stem cells, through TA cells, into secretory and absorptive cells. Fibroblasts enable WT epithelia to access revCSC while retaining secretory and absorptive differentiation. In contrast, *shApc* and *Kras*^*G12D/+*^ limit differentiation and trap cells in the proCSC state. **B)** Data-driven signalling models underpinning the transition from revCSC to proCSC. Arrow colour indicates pathway activation (black on, grey off), while arrow weight depicts relative signalling flux.

When WT colonic epithelia are projected onto this embedding, stem cells occupy high positions in the landscape, with TA cells descending into a central valley before diverging into terminally differentiated secretory and absorptive cells. When WT epithelia communicate with fibroblasts, the TA valley erodes as cells access revCSC. In contrast, CRC mutations *shApc* and *Kras*^*G12D/+*^ resculpt the entire landscape, trapping most cells in the proCSC fate by restricting their differentiation potential (Figure 6A).

The functional perturbation experiments described in this study support a signalling model that underpins each landscape (Figure 6B). In homeostatic WT epithelia, WNT3A, EREG, and R-Spondin-1 drive balanced β-catenin, MAPK, PI3K, and YAP signalling to enable an equilibrium of stem and terminally differentiated cell-fates. When exposed to fibroblast-derived TGF-β1, WT cells become dominated by the YAP signalling flux, have minimal MAPK and PI3K activity, and are therefore polarised towards revCSC. By contrast, APC-loss and KRAS^G12D^ hyper-activate cell-intrinsic β-catenin, MAPK, and PI3K signalling, while simultaneously downregulating receptor expression to decouple epithelia from cell-extrinsic regulation. This limits CRC access to revCSC and traps cells in the proCSC fate. CRC cells can only escape proCSC through high TGF-β1 and low PI3K – tipping the signalling balance back towards revCSC. These observations demonstrate that colonic epithelia exist on an integrated differentiation landscape that can be traversed by co-regulating core signalling hubs, either through cell-intrinsic mutations or cell-extrinsic ligands.

## Discussion

Single-cell technologies can describe cell-type-specific regulation of differentiation and cell-cell communication [31, 32, 33]. In this study, we utilised both multiplexed scRNA-seq and high-throughput MC to functionally map how oncogenic mutations and stromal cues co-regulate colonic epithelia across a continuous polarisation landscape. By analysing >1,000 organoid cultures at single-cell resolution, we identify a stepwise cell-fate trajectory spanning from fibroblast-induced revCSC through an equilibrium of balanced differentiation to oncogene-driven proCSC. While scRNA-seq provides in-depth description of colonic epithelial differentiation and proCSC/revCSC polarisation, multiplexed TOB*is* MC allows comprehensive functional interrogation of cell-intrinsic and -extrinsic cues regulating each cell-fate.

The intestinal stroma comprises a heterogenous population of fibroblasts that regulate the intestinal stem cell niche [2]. In the colonic epithelium, CD34^hi^ fibroblasts located at the crypt bottom are a major source of WNT2B, GREM1, and R-Spondin-1, contributing to both homeostatic stem cell maintenance and tissue regeneration following injury [19]. In contrast, CD34^lo^ fibroblasts reside around upper crypts, show lower expression of WNT2B/GREM1 but higher expression of BMPs, thereby providing a permissive environment for epithelial differentiation [7, 20]. The fibroblasts used in this study contain both CD34^hi^ and CD34^lo^ cells – mimicking *in vivo* heterogeneity (Figure 1B). Both CD34^hi^ and CD34^lo^ fibroblast subpopulations showed comparable polarisation of revCSC (Figure S1B), suggesting the stromal-epithelial communication in organoid co-cultures may be dominated by TGF-β1 signalling (Figure 6B). While this study uses healthy colonic fibroblasts to model homeostatic signalling, it is possible cancer associated fibroblasts (CAFs) will communicate differently with epithelial cells, particularly in CRC. Future cell-cell communication studies between CAF sub-types [34] and defined epithelial genotypes could uncover exceptions to the signalling models described here and therefore provide novel avenues for therapeutic intervention in CRC.

WNT3A is considered a canonical WNT ligand that activates APC/β-catenin signalling, promotes cell proliferation, and reinforces stem cell identity in the intestinal epithelium [35]. It is therefore widely used in colonic organoid culture to compensate for the absence of Paneth cell-derived WNT3A compared with the small intestine [36]. Surprisingly, we found WNT3A alone polarised WT epithelia towards the slow-cycling revCSC fate. Moreover, *shApc* cannot induce revCSC cell-autonomously, indicating revCSC is not immediately downstream of canonical APC/β-catenin signalling (Figure S3A). Our data suggests that WNT3A drives the polarisation to revCSC via YAP (not β-catenin) (Figure 5G), and homeostatic differentiation requires balanced EGF and WNT3A signalling (Figure S4E-F). WT organoids cultured with WENR ligands are enriched for both proCSC and revCSC while depleted of secretory cells and enterocytes (Figures 1H, S2B). Collectively, these observations confirmed that organoid cell-fates can be fine-tuned via competing signalling pathways and organoid culture media should be carefully considered when modelling cell-types of interest (Figures 3A, S4C-F).

proCSC are enriched in CRC organoids and are transcriptionally similar to cells found in human and mouse CRC (Figure S2A). However, we demonstrated that proCSC are also present in WT epithelia and highly enriched in WT organoids cultured with WENR ligands. We therefore do not consider proCSC to be cancer stem cells. Rather than establishing an entirely new cancer-specific cell-fate, our study suggests that oncogenic mutations cell-intrinsically polarise cells to an extreme yet pre-existing proCSC state, while simultaneously disrupting cell-extrinsic regulation of plasticity – trapping cells as proCSC. These results describe cancer as a chronic, unidirectional shift in de-differentiation.

This study charts a continuous polarisation trajectory between revCSC and proCSC in colonic epithelia. In the healthy small intestine, revival stem cells have been demonstrated to act as multipotent stem cells that can be mobilised to replenish traditional LGR5^+^ stem cells in response to tissue damage [7]. Small intestinal revival stem cells are found in the homeostatic small intestine *in vivo* [8, 33] and resemble an early ‘foetal’ stem cell-fate [24, 25]. Here we show that in colonic epithelia, revCSC are enriched by fibroblast-derived WNT3A and TGF-β via epithelial YAP, but only in the context of low PI3K and MAPK signalling. Our work and others now collectively suggest that fibroblasts are master regulators of revival stem cells in both the small intestine and colon.

Although revCSC are most easily accessible in WT epithelia, multiple studies have suggested revCSC also have an important role in CRC [9]. revCSC are candidates for early tumour initiating cells [8] and may confer WNT-inhibitor resistance in CRC [37]. A recent study in human CRC organoids also demonstrated that cancer cells can escape chemotherapy by adopting a slow-proliferating Mex3a^+^ state driven by a low-EGF and high TGF-β culture environment [10]. Our results confirmed that TGF-β can induce revCSC-like cells in CRC organoids, but this process is rare (Figure S3C) and requires low PI3K signalling (Figure 5F). Moreover, we recently demonstrated that cancer associated fibroblasts (CAFs) can also induce a revCSC-like state in CRC patient-derived organoids (PDOs) that protects CRC cells from chemotherapies including fluorouracil, oxaliplatin, and irinotecan [38]. In this model, CAF-chemoprotection can also be overcome by inhibiting YAP signalling – further demonstrating the central role of YAP in revCSC identity. However, CAF-chemoprotection is highly patient-specific, indicating only certain cell-states can be polarised to revCSC in CRC. Collectively, our results and others suggest fibroblast-induced revCSCs may represent an important ‘drug-tolerant persister’ (DTP) state in CRC. Given that targeting cell-plasticity is an emerging area of cancer therapies [39], future studies could target CRC DTP cells by combining YAP inhibitors (to block access to DTP revCSC) with standard chemotherapies (to kill proCSC).

In summary, through single-cell perturbation analysis of >1,000 organoid cultures, we charted a continuous landscape of cell-intrinsic and -extrinsic regulation of colonic stem cell polarisation. We found that colonic stem cell polarity is regulated by competing YAP and PI3K signalling flux, with stromal TGF-β pushing epithelia towards revCSC and CRC mutations trapping epithelia as proCSC. We conclude that cell-fate plasticity is a hallmark of colonic oncogenesis, and that cells can rapidly traverse the colonic differentiation landscape via combinations of oncogenic and stromal signalling.

## Methods

### Colonic Organoid Culture

Wild-type murine colonic organoids and CRC organoids carrying oncogenic mutations (*shApc* (A), *Kras*^*G12D/+*^ (K), *shApc* and *Kras*^*G12D/+*^ (AK), *Kras*^*G12D/+*^ and *Trp53*^*R172H/–*^ (KP), and *shApc, Kras*^*G12D/+*^ and *Trp53*^*R172H/–*^ (AKP)) were a kind gift from Lukas Dow (Cornell University) [40]. *shApc* was induced by Doxycycline treatment at 1 µg mL^−1^ and the efficiency of *Apc* knockdown was monitored with EGFP expression. Organoid base medium was made up of advanced DMEM/F-12 (Thermo 12634010) supplemented with 2 mM l-glutamine (Thermo 25030081), 1 mM N-acetyl-l-cysteine (Sigma A9165), 10 mM HEPES (Sigma H3375), 1 *×* B-27 Supplement (Thermo 17504044), 1 *×* N-2 Supplement (Thermo 17502048), and 1*×* HyClone Penicillin Streptomycin Solution (Fisher SV30010). Colonic organoids were cultured in organoid base medium further supplemented with 100 ng mL^−1^ murine WNT3A (mWNT3A, Peprotech 315-20), 50 ng mL^−1^ mEGF (Thermo PMG8041), 50 ng mL^−1^ mNoggin (Peprotech 250-38), 500 ng mL^−1^ mR-Spondin-1 (Peprotech 315-32), and 10 mM nicotinamide (Sigma N0636). WENR ligands were excluded from all experimental conditions throughout this study unless otherwise stated to ensure cell-cell signalling was not dominated by exogenous recombinant proteins.

For the WENR permutation experiment (Figures 2, 3, S3, and S4), colonic organoids were starved of mWNT3A, mEGF, mNoggin, and mR-Spondin-1 (WENR) for 6 h, split at a ratio of 1:3 (WT, A) or 1:6 (K, KP, AK, and AKP), and seeded as monocultures or fibroblast co-cultures at 5,000 fibroblasts per µL of Matrigel. The cultures were incubated with organoid base medium supplemented with 1 *×* Insulin-Transferrin-Selenium-Sodium Pyruvate (ITS-A) (Thermo 51300044) and 10 mM nicotinamide (Sigma N0636) in addition to the combinations of mWNT3A (100 ng mL^−1^), mEGF (50 ng mL^−1^), mNoggin (50 ng mL^−1^), and mR-Spondin-1 (500 ng mL^−1^) as described in Figure 2. The cells were cultured for 48 h prior to TOB*is* MC analysis (see below).

For the WNT-EGF competition experiment (Figure S4C-F), WT colonic organoids were starved of mWNT3A, mEGF, mNoggin, and mR-Spondin-1 (WENR) for 6 h and split at a ratio of 1:3 and seeded as monocultures. WNT3A ranged from 0 to 100 ng mL^−1^ (0, 10, 20, 50, 100 ng mL^−1^) and / or EGF ranged from 0 to 50 ng mL^−1^ (0, 10, 25, 40, 50 ng mL^−1^) were added to the culture to capture their differential polarisation of revCSC and proCSC. The cells were cultured for 48 h prior to TOB*is* MC analysis (see below).

For the CellChat follow-up experiment (Figures 4D, S5D-F), colonic organoids were starved of mWNT3A, mEGF, mNoggin, and mR-Spondin-1 (WENR) for 6 h, split at a ratio of 1:3 (WT, A) or 1:6 (K, KP, AK, and AKP), and seeded as monocultures. The cells were incubated with organoid base medium supplemented with 1 *×* ITS-A (Thermo 51300044), 10 mM nicotinamide (Sigma N0636), and the signalling ligands identified from the ligand-receptor analysis (Figure 4C): murine WNT5A (250 ng mL^−1^, R&D Systems 645-WN-010/CF), murine SEMA3A (250 ng mL^−1^, R&D Systems 5926-S3-025/CF), human TGF-β2 (1 ng mL^−1^, BioLegend 583301), murine TGF-β1 (1 ng mL^−1^, BioLegend 763102), murine IGF1 (100 ng mL^−1^, Cell Guidance Systems GFM5-10), murine NRG1 (100 ng mL^−1^, R&D Systems 9875-NR-050), murine EREG (500 ng mL^−1^, R&D Systems 1068-EP-050/CF), and murine OPN (400 ng mL^−1^, BioLegend 763604). Organoids treated with WNT3A (100 ng mL^−1^) or EGF (50 ng mL^−1^) were included as positive controls. The cells were cultured for 48 h prior to TOB*is* MC analysis (see below).

For the cue-signal-response MC array (Figures 5, S6), colonic organoids were starved of mWNT3A, mEGF, mNoggin, and mR-Spondin-1 (WENR) for 6 h, split at a ratio of 1:3 (WT, A) or 1:6 (K, AK), and seeded as monocultures. The cells were incubated with organoid base medium supplemented with 1 *×* ITS-A, 10 mM nicotinamide, with or without signalling ligands: murine WNT3A (100 ng mL^−1^), murine EREG (500 ng mL^−1^), murine TGF-β1 (2 ng mL^−1^). For each ligand condition, signalling inhibitors were added at the following concentrations: CA3 (YAP inhibitor, 2 µM, Sigma SML2647), CHIR99021 (GSK-3β inhibitor, 3 µM, Cell Guidance Systems SM13-1), ICG-001 (CBP/β-Catenin inhibitor, 2 µM, Cayman Chemical 16257), Dasatinib (SRC inhibitor, 50 nM, Cell Guidance Systems SM45-20), Trametinib (MEK inhibitor, 50 nM, Cayman Chemical 16292), GDC-0941 (PI3K inhibitor, 1 µM, Selleck Chemical 50-851-6), PF-573228 (FAK inhibitor, 2.5 µM, Cayman Chemical CAY14924), and SIS3 (SMAD3 inhibitor, 3 µM, Cayman Chemical 15945). The cells were cultured for 48 h prior to TOB*is* MC analysis (see below).

### Heterocellular Organoid Culture

The heterocellular organoid cultures were established as previously described [12]. Briefly, organoids were starved of mWNT3A, mEGF, mNoggin, and mR-Spondin-1 (WENR) for 6 h prior to the experiment and passaged at a ratio of 1:2.5; colonic fibroblasts (isolated, immortalised, and characterised in [12]) were seeded at 6,000 cells per µL for monoculture, 5,000 cells per µL for two-way co-cultures, and 4,000 cells per µL for three-way co-cultures; primary bone marrow-derived macrophages were seeded at 9,000 cells per µL for monoculture, 8,000 cells per µL for two-way co-cultures, and 7,000 cells per µL for three-way co-cultures. The cells were mixed in Matrigel and seeded at 7 *×* 40 µL droplets per well in 6-well plates (for scRNA-seq) or

1 *×* 50 µL droplet per well in 48-well plates (for TOB*is* MC). Unless otherwise specified, each microenvironment culture was maintained in WENR-free advanced DMEM/F-12 (Thermo 12634010) supplemented with 2 mM l-glutamine (Thermo 25030081), 1 mM N-acetyl-l-cysteine (Sigma A9165), 10 mM HEPES (Sigma H3375), 1 *×* B-27 Supplement (Thermo 17504044), 1 *×* N-2 Supplement (Thermo 17502048), 1 *×* Insulin-Transferrin-Selenium-Sodium Pyruvate (ITS-A, Thermo 51300044) and 1 *×* HyClone penicillin streptomycin solution (Fisher SV30010) for 48 h prior to TOB*is* MC analysis (see below).

### scRNA-seq Data Acquisition

To prepare single-cell suspensions from the heterocelluar organoid cultures, cells were removed from Matrigel using ice-cold PBS, collected with a benchtop centrifuge, and incubated with TrypLE™ Express Enzyme (Thermo 12604013) for 7 to 10 min at 37 °C. The cells were then washed with ice-cold advanced DMEM/F-12 (Thermo 12634010) and filtered through a 35-µm cell strainer (Fisher 10585801). For FACS sorting, eBioscience™ Fixable Viability Dye eFluor™ 780 (FVD780, Thermo 65-0865-14) was used to label dead cells, while FITC anti-mouse CD66a (CEACAM1) antibody (Clone: MAb-CC1; BioLegend 134518) was used to stain epithelial cells, and APC anti-mouse CD45 antibody (Clone: BM8; BioLegend 123116) was used to stain macrophages. The gating of fibroblasts was based on their endogenous DsRed expression [12]. The collected cells were counted with a Countess II automated cell counter (Thermo Fisher) and examined for viability (samples with >90% viable cells were passed onto scRNA-seq library construction). To preserve RNA in the samples and to minimise technical variations, cells were fixed in ice-cold methanol immediately after counting as per the 10X Genomics instruction. For co-cultures, different cell-types were mixed at equal cell numbers prior to the fixation step. The methanol-fixed cells were stored at −20 °C for up to 2 weeks before they were rehydrated and processed using the 10X Genomics Chromium Controller. scRNA-seq libraries were generated with the 10X Genomics Chromium Next GEM Single Cell 3’ Reagent Kits v3.1 (Dual Index) and sequenced with the Illumina NovaSeq 6000 System (2 *×* 150 bp paired-end reads), aiming at 60,000 read pairs per cell and 2,000 cells per cell-type per sample.

### scRNA-seq Data Processing

Raw binary base call (BCL) sequence files were converted to FASTQ files and processed with the 10X Genomics Cell Ranger pipeline version 5.0.1. The FASTQ files were then aligned to a custom GRCm38 reference genome containing the sequences of *DsRed* and *eGFP* transgenes present in fibroblasts and organoids respectively, generating pre-filtered feature-barcode matrices.

The gene count matrices were analysed with the R package *Seurat* version 4.0.4 [16]. The analysis pipeline encompasses quality control, data normalisation, data integration, dimensionality reduction, cell clustering, and analysis of differential gene expression. Genes found in less than 4 cells were removed during QC and only cells with > 600 unique genes identified were kept for downstream analysis. The total number of detected sequences typically ranged from 1,200 to 80,000 per cell, and the actual values were manually determined based on cell-type composition and sequencing depth. For the integrated epithelial object in Figure 1D, an additional filtering step was performed to remove cells with undetectable expression for any one of the bona fide pan-epithelial genes *Epcam, Krt8, Krt18, Krt19, Cldn7*. Cell-cycle regression was performed using the *sctransform* function. Log-normalised gene expression values (RNA assay) were used for downstream analysis if not otherwise stated.

Dataset integration was performed using Seurat’s reciprocal PCA (RPCA) implementation [16] (k.anchor=12) as it has been optimised to handle large datasets. The integrated object in Figure 1B was computed using all cells from the 20 conditions shown in Figure 1A (integrated object limited to 2,000 genes across 58,726 cells). The integrated object in Figure 1D was computed using just the epithelial cells from all conditions (4,000 genes, 29,452 cells).

For dimensionality reduction (DR), the first 50 principal components (PC) was computed from the integrated assays to generate 2-dimensional PHATE embeddings with default parameters (Table S2). PHATE was chosen as the standard DR method for the study due to its capacity to capture the global structure in biological systems with important developmental trajectories [17].

Cell clustering was computed using the Leiden algorithm on the *k* NN graph generated from the integrated epithelial dataset (first 48 PCs), at a series of resolutions ranging from 0.2 to 0.8. The final cluster annotations were retrospectively defined by common cell-type marker expression, inter-cluster relationships on a multi-resolution clustering tree [41], and cross-condition differential abundance behaviours (see below). Cells from outlier clusters (totalling less than 1% of all epithelial cells) were excluded from the downstream analysis.

Differentially expressed (DE) genes between clusters, conditions, and cell neighbourhoods were identified using Wilcoxon rank-sum test implemented by Seurat’s *FindAllMarkers* and *FindMarkers* functions.

### scRNA-seq Data Analysis

To generate the EMD-PCAs in Figure 1C, log-normalised gene expression data of all cells of a particular cell-type (epithelial cells, fibroblasts, or macrophages) were exported from the integrated object. EMD scores for the top 6,000 variable genes of each condition were calculated with CyGNAL [42] using the WT monoculture control as the reference.

Differentially abundant (DA) cell neighbourhoods were identified using the R package *MiloR* [23], which enabled the detection of enrichment and depletion of cell clusters caused by microenvironmental and/or genotypical perturbations. Given that CD34^hi^ and CD34^lo^ fibroblasts do not differentially regulate epithelial cells (Figure S1B), all samples of WT organoid+fibroblast co-cultures were grouped and considered replicates of the query condition regardless of the CD34 status of the fibroblasts, with the DA test threshold set at 5% SpatialFDR (Figure 1F). Similarly, AK and AKP organoid monocultures were grouped due to their similar DE and DA behaviour (Figure 1G). The DA overview dot plot in Figure 1H was generated by comparing the 17 conditions against the WT monoculture control (2*×* replicates).

Heatmaps of selected marker genes were generated with the R package *ComplexHeatmap* [43] across the manuscript. Gene lists in Figures 1I and Figure S1B were curated from previously reported markers for colonic epithelial subpopulations and DE genes detected between epithelial clusters, conditions, and DA neighbourhoods within this study. Gene lists in For S1B-D represent DE genes between conditions.

The *UCell* [44] method was used to generate the correlation matrix between gene signatures in existing literature and cell clusters identified within this study (Figure S2A). Gene lists for different intestinal stem cell-states were compiled from public datasets, together with transcriptional targets of key signalling pathways encoding the different stem cell-states (Table S3). These gene lists were compared with the curated gene signatures for proliferation, CSC, revCSC, and proCSC cell-states in Figure 1I, as well as the top DE genes for each stem cluster (adjusted p-value < 0.01, log2FC > 0.25, top 24 genes with the greatest positive log2FC values) (Table S3). *UCell* scores for each gene set were calculated using Log-normalised gene expression values and z-scored to allow cross-signature comparison. Pearson correlations were computed between the scores on all cells of stem and TA clusters and then visualised as a correlation heatmap, grouped via complete linkage hierarchical clustering, only showing significant correlations (conf.level = 0.95).

Leveraging the concept that cells with a higher potency should have a higher signalling entropy [45], the pluripotency values for epithelial cells across the different clusters were estimated using the R package *SCENT* [21]. Signalling entropy scores for all epithelial cells were computed with the CCAT (correlation of connectome and transcriptome) approximation method using a murinised version of the built-in *net17Jan16* Protein-Protein interaction network.

For RNA velocity analysis, loom files were generated from Cell Ranger’s output using the Python package *velocyto* [46] (reference genome: GRCm38, repeat mask assembly: GRCm38/mm10, track: RepeatMasker). RNA velocity was analysed with the Python package *scVelo* [22] using default parameters unless otherwise specified (Table S2). Metadata and PHATE embedding coordinates were exported from the relevant Seurat objects to filter and annotate anndata objects generated from the loom files made by velocyto. Moments for the velocity estimation were calculated using the first 50 PCs and 30 neighbours from the anndata objects. RNA velocities were computed with the *recover_dynamics* function using the dynamical model of transcriptional dynamics with default parameters. The velocity stream embedding (Figure 1E) was computed using the integrated object containing epithelial cells from all conditions. The RNA velocity vector lengths, an estimate of a cell’s differentiation rate, were computed using cells solely from the 4 conditions shown in Figure S2B-D. The quantitative comparison in Figure S2D was performed using the Games-Howell pairwise test wrapper from the R package *statsExpressions* [47]. All conditions were compared against the WT monoculture control and all *p*-values have been corrected for multiplicity with the Holm method.

Ligand-receptor expression analysis was performed using the R package CellChat [28], where stromal-epithelial signalling was analysed across 4 different organoid genotypes (WT, A, AK, and AKP). Epithelial cells were annotated with the clusters previously identified (Figure 1D), while the fibroblasts were grouped as a single cluster. A merged CellChat object was generated to compare relative communication probability of fibroblast-to-epithelia signalling across the genotypes. Significant ligand-receptor pairs were identified based on CellChat’s murine cell communication database. Plots displaying aggregate outgoing and incoming communication probability (Figure 4A) were generated with the *netAnalysis_-signalingRole_scatter* function. Detected communication at the pathway and interaction level was accessed with the *subsetCommunication* function and probabilities were z-score normalised to allow for cross-pathway or cross-interaction comparison. The results were visualised with ComplexHeatmap in Figure 4B-C, the rows of which were manually ordered based on hierarchical clustering and grouped based on the nature of the interaction. Gene expression of the ligand-receptor pairs identified above was visualised using Seurat’s *Dotplot* function in Figure S5A. *UCell* scores for ligand and receptor genes were calculated for fibroblasts and epithelial cells respectively. Games-Howell pairwise test was performed using the R package *statsExpressions* and all *p*-values have been corrected for multiplicity with the Holm method.

### TOB*is* MC

TOB*is* MC of organoid cultures was performed as previously described [13]. Briefly, the cultures were incubated with 25 µM ^127^5-iodo-2’-deoxyuridine (^127^IdU) for 30 min to label S-phase cells, treated with a cocktail of protease (Sigma P8340) and phosphatase inhibitors (Sigma 4906845001) to protect protein and phosphorylation epitopes, and fixed with 4% (w/v) PFA for 1 h at 37 °C. The cells were washed twice with PBS, incubated in 250 nM ^194/8^cisplatin (Fluidigm 201194/8) for 10 min to stain dead cells, and washed twice with PBS to remove residual cisplatin. TOB*is* barcodes were added to the cells and incubated overnight at 4 °C. The following day, unbound barcodes were quenched with reduced glutathione (Sigma G6529) and washed from the cultures. TOB*is*-barcoded organoids from each condition were removed from Matrigel in a freshly prepared dissociation buffer containing 0.5 mg mL^−1^ Dispase II (Thermo 17105041), 0.2 mg mL^−1^ Collagenase IV (Thermo 17104019) and 0.2 mg mL^−1^ DNase I (Sigma DN25), pooled into a single master tube and dissociated into single cells with a gentleMACS Octo Dissociator (Miltenyi 130-096-427). Following dissociation, the cells were washed, filtered, and stained for extracellular epitopes with rare earth metal-labelled antibodies (Table S1). The cells were then permeabilised with 0.1% (v/v) Triton X-100 followed by 50% (v/v) methanol. Once permeabilised, the cells were stained with a panel of metal antibodies against intracellular proteins and PTMs (Table S1). For each cell, we measured cell-type markers (epithelia: CEACAM-1, Pan-cytokeratin (Pan-CK), GFP; fibroblasts: PDPN, RFP, mCherry), epithelial differentiation markers identified by scRNA-seq (CLU, CD44, SOX9, SURVIVIN, LRIG1, EPHB2, C-MYC, and FABP2), cell-state markers (pRB [S807/S811], IdU, pHH3 [S28], Cyclin B1, and cCaspase3 [D175] [13]), and >20 PTMs spanning multiple cell-signalling pathways. The cells were washed and incubated in DNA intercalator ^191/193^Ir (Fluidigm 201192A) overnight before MC single-cell data acquisition and analysis.

### MC Data Acquisition and Analysis

TOB*is* MC data were acquired and analysed as previously described [13]. For Fluidigm Helios acquisitions, stained cells were washed into Maxpar Water (Fluidigm 201069) containing 2 mM EDTA, diluted to 0.8–1.2 *×* 10^6^ cells mL^−1^ and spiked with EQ Four Element Calibration Beads (Fluidigm 201078). The cells were then loaded into a Super Sampler (Victorian Airships). For CyTOF XT acquisitions, stained cells were wash into Maxpar Cell Acquisition Solution Plus (Fluidigm 201244) containing 2 mM EDTA, diluted to 0.8–1.2 *×* 10^6^ cells mL^−1^ and spiked with EQ™ Six Element Calibration Beads (Fluidigm 201245).

After data acquisition, raw MC data were normalised and exported as standard FCS file(s). Multiplexed TOB*is* experiments were debarcoded into individual conditions (https://github.com/zunderlab/single-cell-debarcoder), imported into Cytobank (http://www.cytobank.org/), and gated with Gaussian parameters, DNA/cisplatin, and cell-type markers to remove debris, identify live cells, and remove doublets respectively. The fully gated datasets were further processed with our MC data analysis pipeline, CyGNAL (https://github.com/TAPE-Lab/CyGNAL) [42]. Earth mover’s distance (EMD) [48] was used to quantify node intensity of each marker. Unless otherwise specified, EMD scores were calculated with WT untreated controls (concatenated replicates) as the reference.

PHATE [17] embeddings were calculated with raw/z-scored EMD scores or arcsinh-transformed single-cell MC data using the python package *phate* (https://github.com/KrishnaswamyLab/PHATE) with parameters specified in Table S2. EMD heatmaps were generated with the R package ComplexHeatmap [43] and further annotated in OmniGraffle Professional across the manuscript. For the WENR permutation experiment (Figures 2, 3, S3, and S4), EMD scores for revCSC and proCSC markers (Figure S3A), percentages of S-phase cells (Figure S3B) and CLU^+^ cells (Figure S3C) were plotted and analysed with GraphPad Prism 7 (ordinary one-way ANOVA with Holm-Šídák’s multiple comparisons test for Figure S3A, unpaired two-tailed *t* - tests for Figure S3B-C). For the cue-signal-response perturbation array (Figures 5, S6), EMD scores for CLU and LRIG1 were calculated for selected conditions and analysed with GraphPad Prism 7 (ordinary one-way ANOVA with Holm-Šídák’s multiple comparisons or unpaired two-tailed *t* -tests for Figure 5G, H).

Force-directed Scaffold Maps [27] (Figure 3C) were constructed using the R package Scaffold (https://github.com/nolanlab/scaffold). Landmark populations (WT, A, K, KP, AK, AKP organoid monocultures) were manually gated and exported from Cytobank with all data arcsinh transformed (cofactor = 5). The parameters used in the Scaffold analysis were specified in Table S2.

The Boolean logic models of CSC regulation by WENR ligands (Figure 3D) were compiled in OmniGraffle Professional.

The relative stemness (RS) score (Figure 5F) was generated by calculating ratios between arcsinh-transformed LRIG1 and CLU MC measurements for single cells, followed by log_2_ normalisation, and then summarised at the replicate and condition level. RS score heatmap (Figure S6B) was generated with the R package ComplexHeatmap [43] and further annotated in OmniGraffle Professional.

### Valley-Ridge (VR) Score

The VR score was defined as the weighted sum of the Valley (weight = 0.9) and the Ridge (weight = 0.1) components, and was computed on a per sample and per cluster basis (Figure S7A). The Valley component equals the median CCAT value of each sample-cluster combination. To calculate the Ridge component, the inverse of the velocities was first computed and scaled to a range between 0 and 1. A cell centrality distance was then calculated for cells in each cluster by first building a *k* NN graph of a cluster’s cells from the PHATE embeddings (Table S2), followed by the calculation of a distance matrix using graph-tool’s *shortest_path* function [49]. The median distance for each cell to all other cells was then calculated, whereby cells with the lowest distance would be at a cluster’s centre whilst those with the highest distance would be at the cluster periphery. To allow intercluster comparisons, outliers with a distance over *Q*_99_ were set to the median distance value before scaling to (0,1). Finally, the Ridge component was computed per sample-cluster as the product of the median scaled inverse velocities and the cell’s scaled centrality distance (Figure S7A).

This definition of the VR score allows the CCAT-driven Valley component to be the driving force for sculpting the landscape and the velocity-driven Ridge component to predominately define the barriers around clusters – producing a tarn-like effect symbolising a state of trapped cells. In principle, any other dimensionality reduction technique can be used in place of PHATE [50], and the Valley/Ridge component can be computed using other metrics underpinning pluripotency and cell-fate transition. The Ridge component can also be calculated with a distance-free approach such as α-shapes [51]. Finally, the VR scores can be computed on a per cell or neighbourhood basis, which will increase landscape resolution and liberate the method from constraints of cluster definitions (at the expense of increased noise).

### Waddington-like Landscape

To generate the Waddington-like landscapes in Figure 6A, we combine the ability of PHATE to capture the global structure of single-cell data with the VR score (described above) (Figure S7A-B).

Waddington-like landscapes can be visualised directly in Python (Figure S7B, C). Briefly, a low dimensional 34×30 mesh grid was generated from the PHATE embeddings, and a 3D surface was rendered by projecting VR scores onto the grid using the radial basis function interpolation from scipy [52] (Table S2). The surface of the landscape was coloured by VR scores and a scatter plot was overlaid where the elevation of each cell was defined as the weighted sum of its VR score (weight = 0.9), CCAT value (weight = 0.1), and a constant factor of 0.012 (weight = 1). This added a level of controlled noise to the scatter plot while ensuring most cells remain above the interpolated surface (Figure S7C).

These landscapes can also be visualised in SideFX Houdini 19.5 (http://www.sidefx.com) and rendered using Maxon Redshift 3.5 (http://www.redshift3d.com) (Figures 6A, S7B). VR scores and scRNA-seq metadata were imported and points were positioned in z- and x-axes according to their PHATE scores. This PHATE distribution was then transformed in the y-axis according to each cell’s VR score. The PHATE-transformed 2D distribution was used as a deformation lattice to influence nearby points on a polygonal grid, and its difference from the VR-transformed 3D distribution was used to drive deformation of this polygonal grid into a Waddington-like landscape. The VR-transformed data was then projected back onto the Waddington-like landscape to avoid intersections between positions of data points and landscape topology. A video tutorial to visualise Waddington-like embeddings using Houdini is available at: https://entagma.com/houdini-tutorialwaddington-landscape/.

## Data Availability

Raw scRNA-seq data and BioSample metadata have been deposited at Sequence Read Archive (SRA) (https://www.ncbi.nlm.nih.gov/bioproject/PRJNA883610). Raw and processed MC data are available as a Community Cytobank project (https://community.cytobank.org/cytobank/experiments#project-id=1460). Aligned scRNA-seq count matrices, spliced/unspliced RNA count matrices, integrated Seurat objects, integrated MC dataframes, and Houdini project files can be accessed at Zenodo (https://doi.org/10.5281/zenodo.7586958). All analysis scripts to reproduce figure plots together with a notebook explaining pre-processing and QC steps for scRNA-seq analysis are available at GitHub (https://github.com/TAPE-Lab/Qin-CardosoRodriguez-et-al).

## Acknowledgements

We are extremely grateful to L. Dow for sharing murine colonic organoids and S. Acton for providing murine tissue for fibroblast and macrophage isolation. We thank Y. Guo and the UCL CI Flow-Core for CyTOF support. We thank E. Sahai, V. Li, S. Acton, and members of the Tape Lab for their constructive critique of the manuscript. This work was supported by Cancer Research UK (C60693/A23783), the Cancer Research UK City of London Centre (C7893/A26233), the UCLH Biomedical Research Centre (BRC422), and the UKRI Medical Research Council (MR/T028270/1).

## Author Contributions

X.Q. designed the study, performed organoid experiments, generated scRNA-seq and TOB*is* mass cytometry data, analysed mass cytometry data, and wrote the paper. F.C.R. analysed scRNA-seq data, developed the VR score, and wrote the paper. J.S. developed TOB*is* barcodes and conjugated rare-earth metal antibodies. P.V. provided organoid culture support. J.C rendered Waddington-like landscapes. C.J.T. designed the study, analysed the data, and wrote the paper. Co-authors reserve the right to rearrange authorship positions on their CVs.

## Supplementary Information

### Supplementary Figures

**Figure S1.**
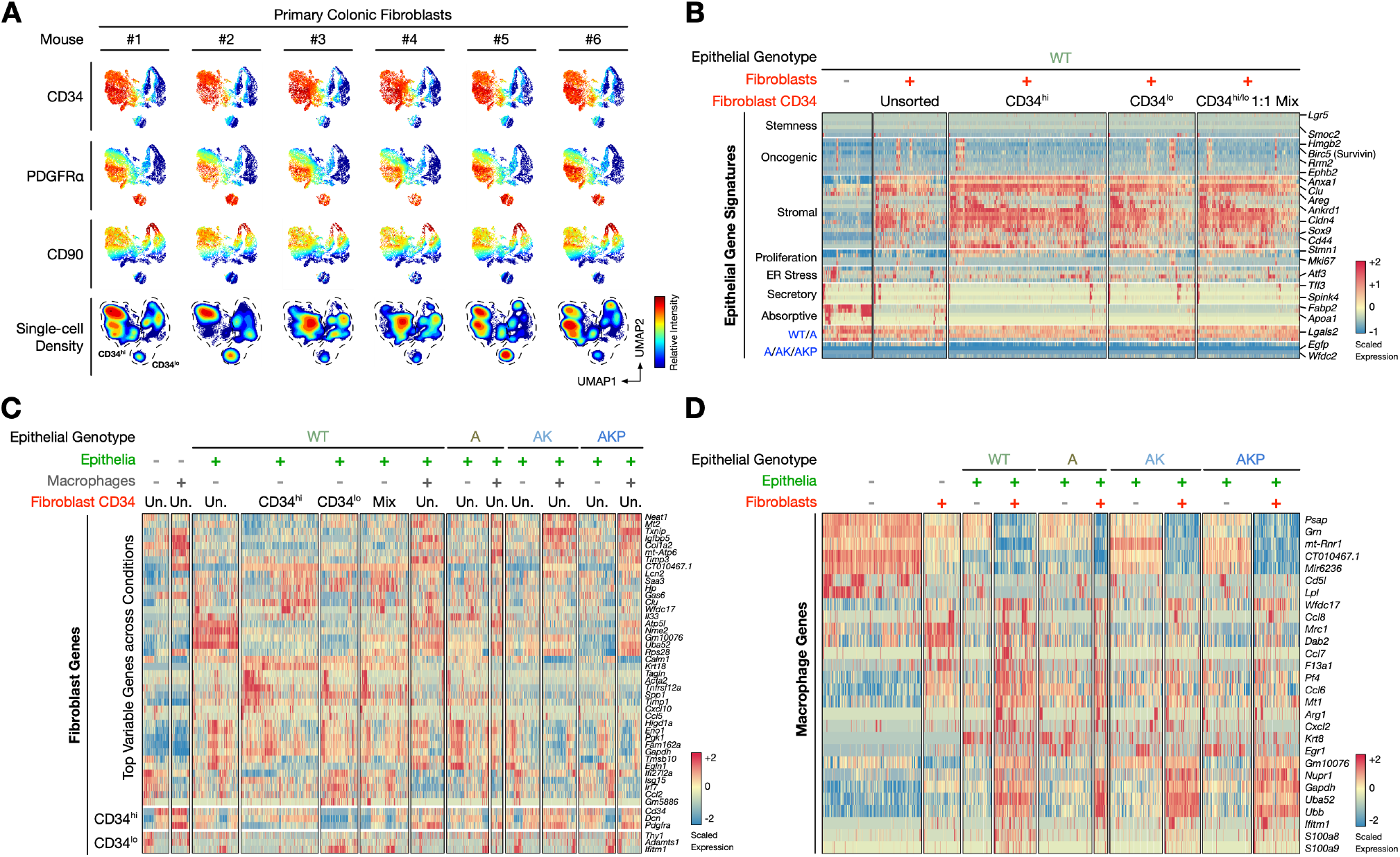
Single-cell Characterisation of the Heterocellular Organoid Model. Related to Figure 1. **A)** Mass cytometry (MC) analysis of primary murine colon fibroblasts showing stromal markers CD34, PDGFRα, and CD90. **B)** scRNA-seq analysis of WT colonic organoids co-cultured with unsorted, CD34^hi^, CD34^lo^, and a 1:1 mix of CD34^hi^:CD34^hi^ colonic fibroblasts. **C)** Differential gene expression analysis of fibroblasts regulated by epithelial organoids and macrophages. **D)** Differential gene expression analysis of macrophages regulated by epithelial organoids and fibroblasts.

**Figure S2.**
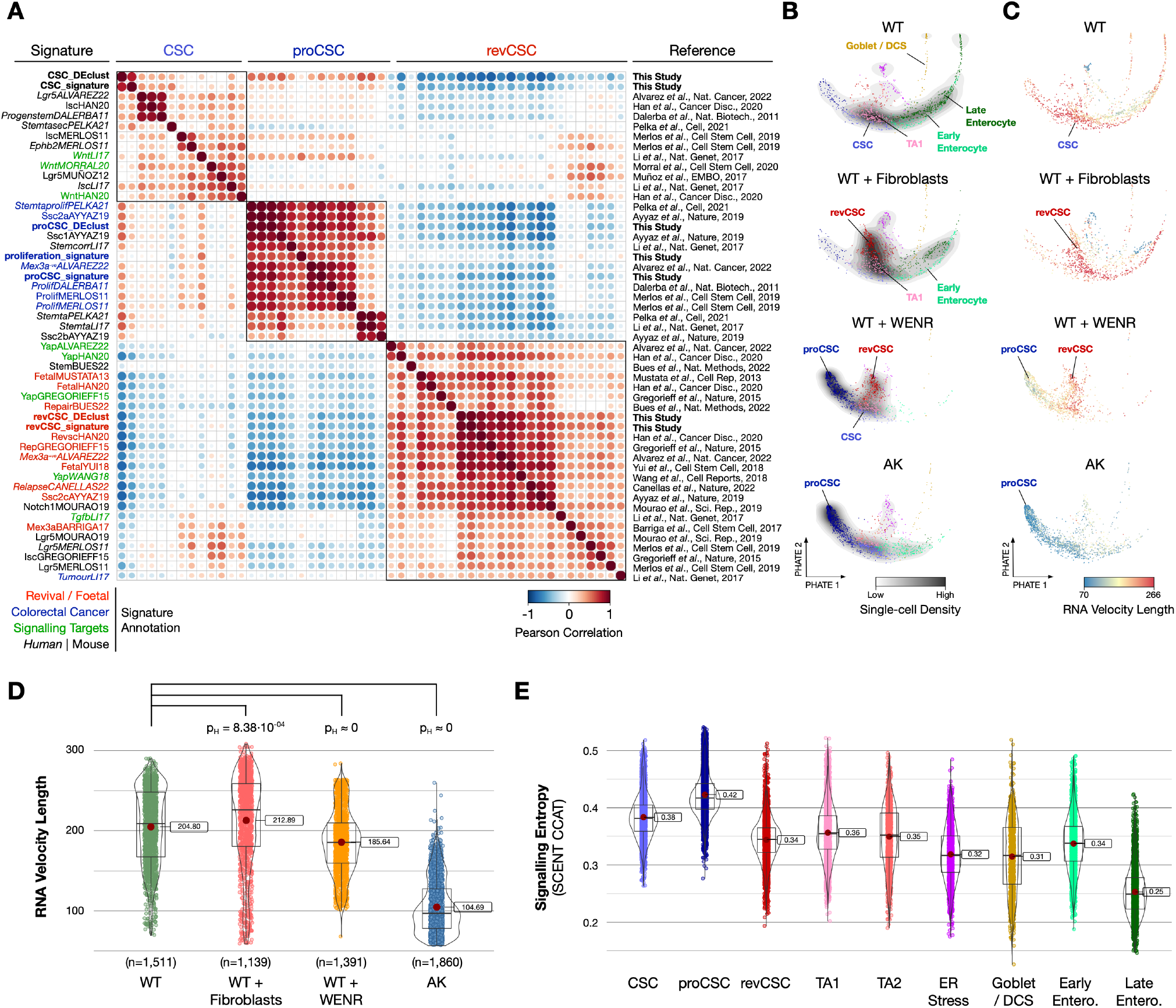
Epithelial Stem Cell Signatures. Related to Figure 1. **A)** Comparison of gene signatures of CSC, proCSC, and revCSC identified in this study with published stem cell signatures. **B)** Single-cell PHATE embeddings of epithelial cells from WT, WT+Fibroblasts, WT+WENR, and AK organoids coloured by cluster and overlaid with single-cell density. **C)** Single-cell PHATE embeddings coloured by RNA velocity vector lengths. **D)** RNA velocity vector lengths of organoid conditions (Games-Howell pairwise test with Holm-adjusted *p*-values). **E)** CCAT scores of epithelial clusters. CSC, colonic stem cell. proCSC, hyper-proliferative CSC. revCSC, revival CSC. TA, transit amplifying cell. DCS, deep crypt secretory cell. Entero., enterocyte. Boxplots show min/max and quartiles. Red dot marks the mean value.

**Figure S3.**
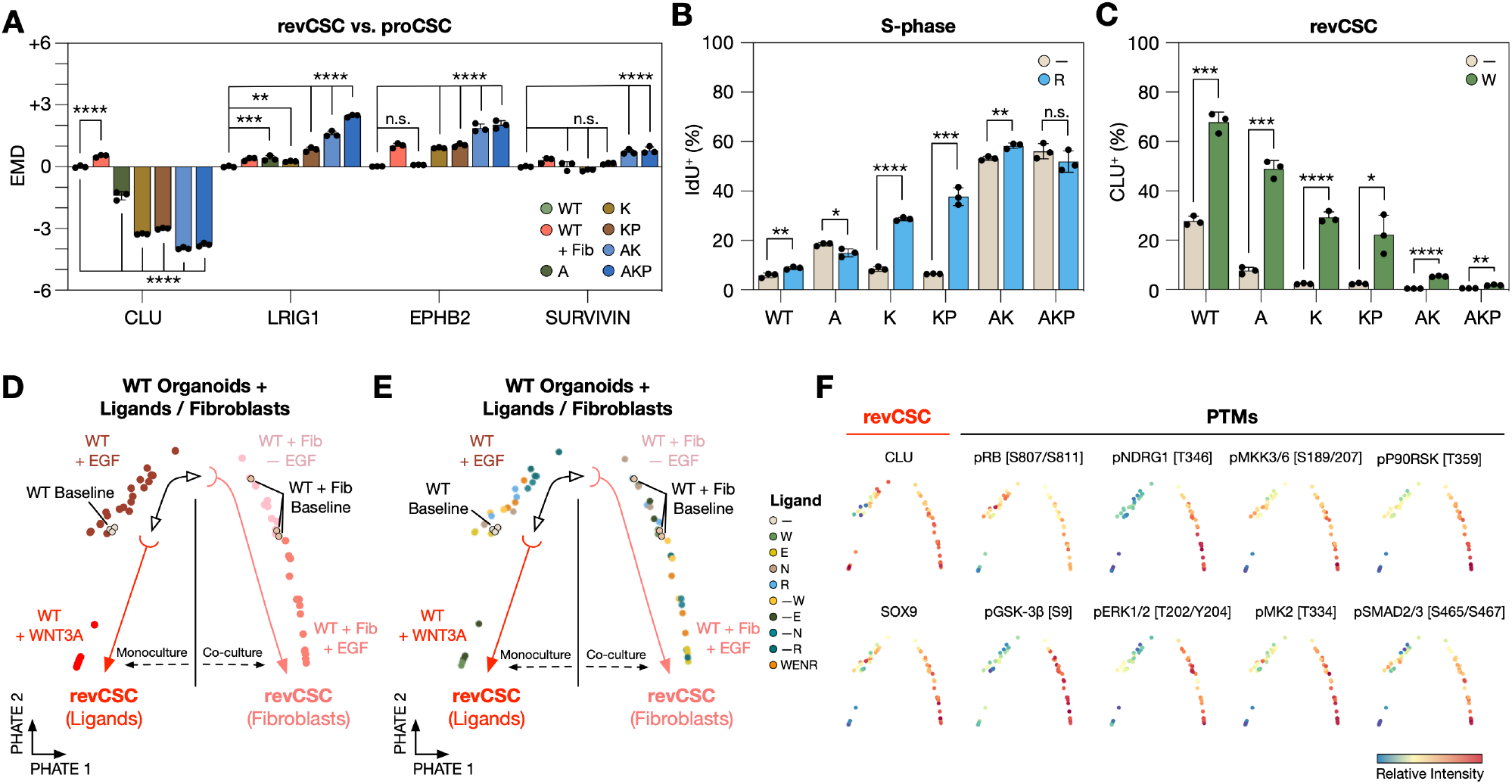
Epithelial Regulation by Organoid Genotypes, WENR Ligands, and Fibroblasts. Related to Figure 2. **A)** EMD scores for CLU, LRIG1, EPHB2, and SURVIVIN across organoid culture conditions (**, *p* < 0.01; ***, *p* < 0.001; ****, *p* < 0.0001. Ordinary one-way ANOVA with Holm-Šídák’s multiple comparisons test). Error bars represent standard deviation (SD). **B)** Percentage of S-phase cells in organoids cultured with or without R-Spondin-1 across genotypes (***, *p* < 0.001; ****, *p* < 0.0001. Unpaired two-tailed *t* -test). Error bars represent SD. **C)** Percentage of revCSC in organoids cultured with or without WNT3A across genotypes (***, *p* < 0.001. Unpaired two-tailed *t* -test). Error bars represent SD. **D-F)** EMD-PHATE of WT organoids cultured with or without fibroblasts and WENR ligands coloured by microenvironment, ligands, and EMD scores for selected markers. One dot = one condition. revCSC, revival colonic stem cell. proCSC, hyper-proliferative colonic stem cell.

**Figure S4.**
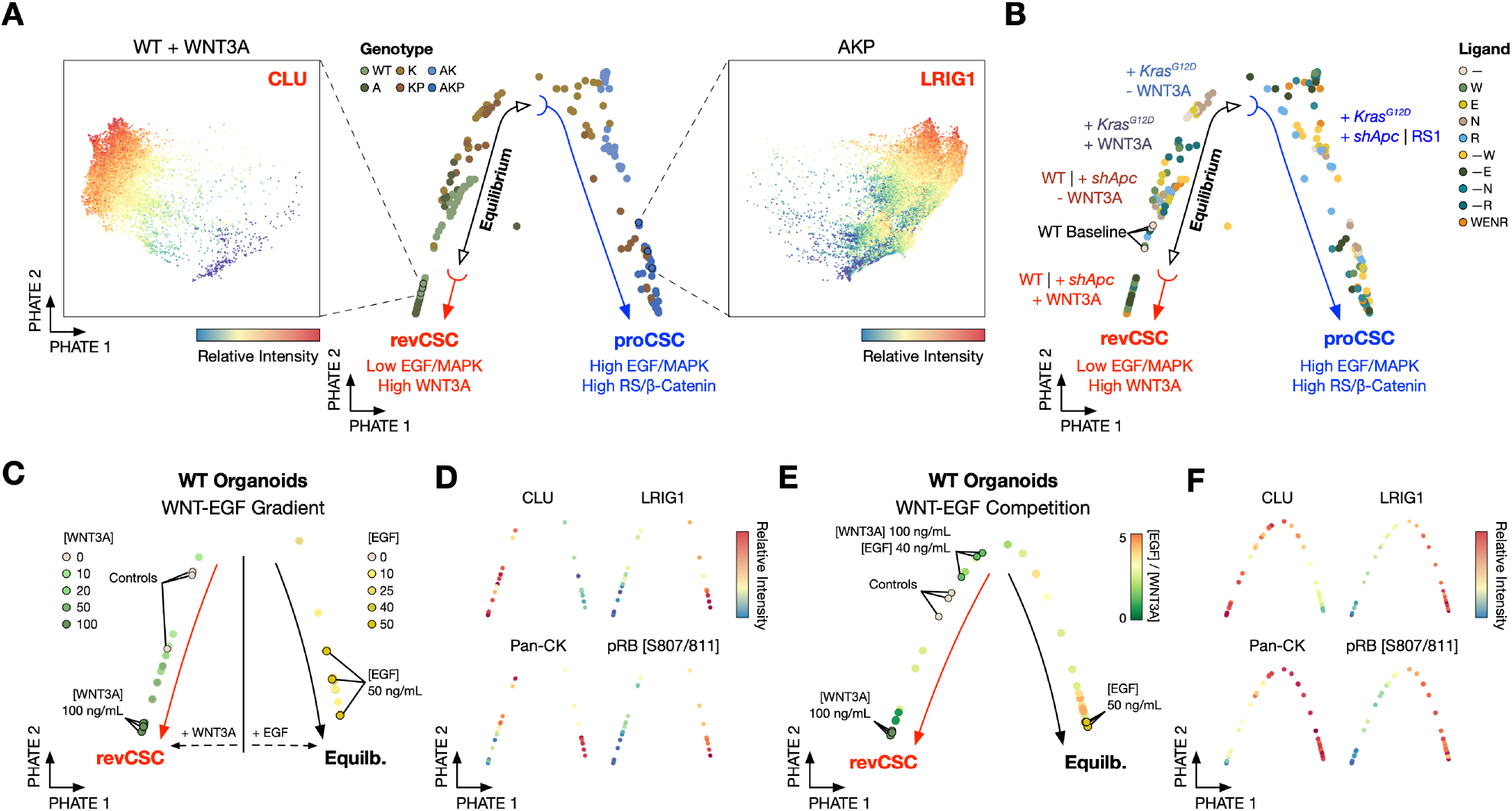
Stepwise Regulation of Epithelial Differentiation. Related to Figure 3. **A)** EMD-PHATE of organoid monocultures +/-WENR ligands coloured by organoid genotypes. Inserts: single-cell PHATE overlaid with arcsinh-transformed measurements of CLU or LRIG1 for WT+WNT3A versus AKP monoculture respectively. **B)** EMD-PHATE of organoid monocultures +/-WENR ligands annotated by organoid genotype and culture conditions. **C)** EMD-PHATE of WT organoids cultured with a gradient of either WNT3A or EGF coloured by WNT3A or EGF concentrations (ng mL^−1^). **D)** The PHATE embedding in **C)** coloured by EMD scores for selected markers. **E)** EMD-PHATE of WT organoids cultured with varying combinations of WNT3A and EGF coloured by the ratio between EGF and WNT3A concentrations. **F)** The PHATE embedding in **E)** coloured by EMD scores for selected markers. Equilb., Equilibrium.

**Figure S5.**
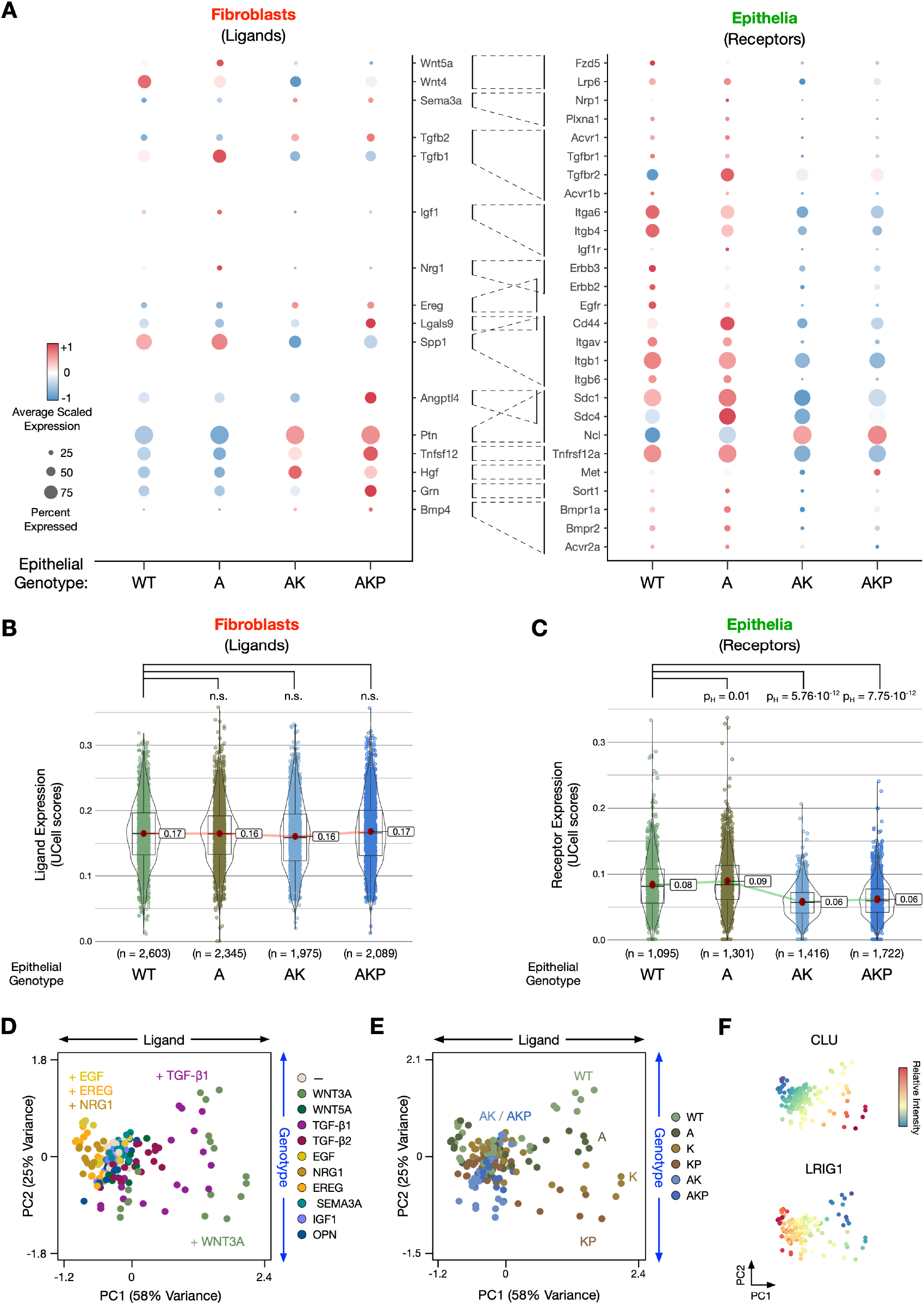
Ligand-receptor Expression Analysis. Related to Figure 4. **A)** Average scaled expression of ligands (expressed by fibroblasts) and receptors (expressed by epithelia) across organoid genotypes. **B)** Ligand expression (*UCell* scores) by fibroblasts in co-cultures across organoid genotypes (Games-Howell pairwise test, n.s not significant). **C)** Receptor expression (*UCell* scores) by epithelia in co-cultures across organoid genotypes (Games-Howell pairwise test with Holm-adjusted *p*-values). **D-E)** EMD-PCA of epithelial cells regulated by exogenous ligands. **F)** PCA from **D)** coloured by EMD scores for CLU and LRIG1. Boxplots show min/max and quartiles. Red dot marks the mean value.

**Figure S6.**
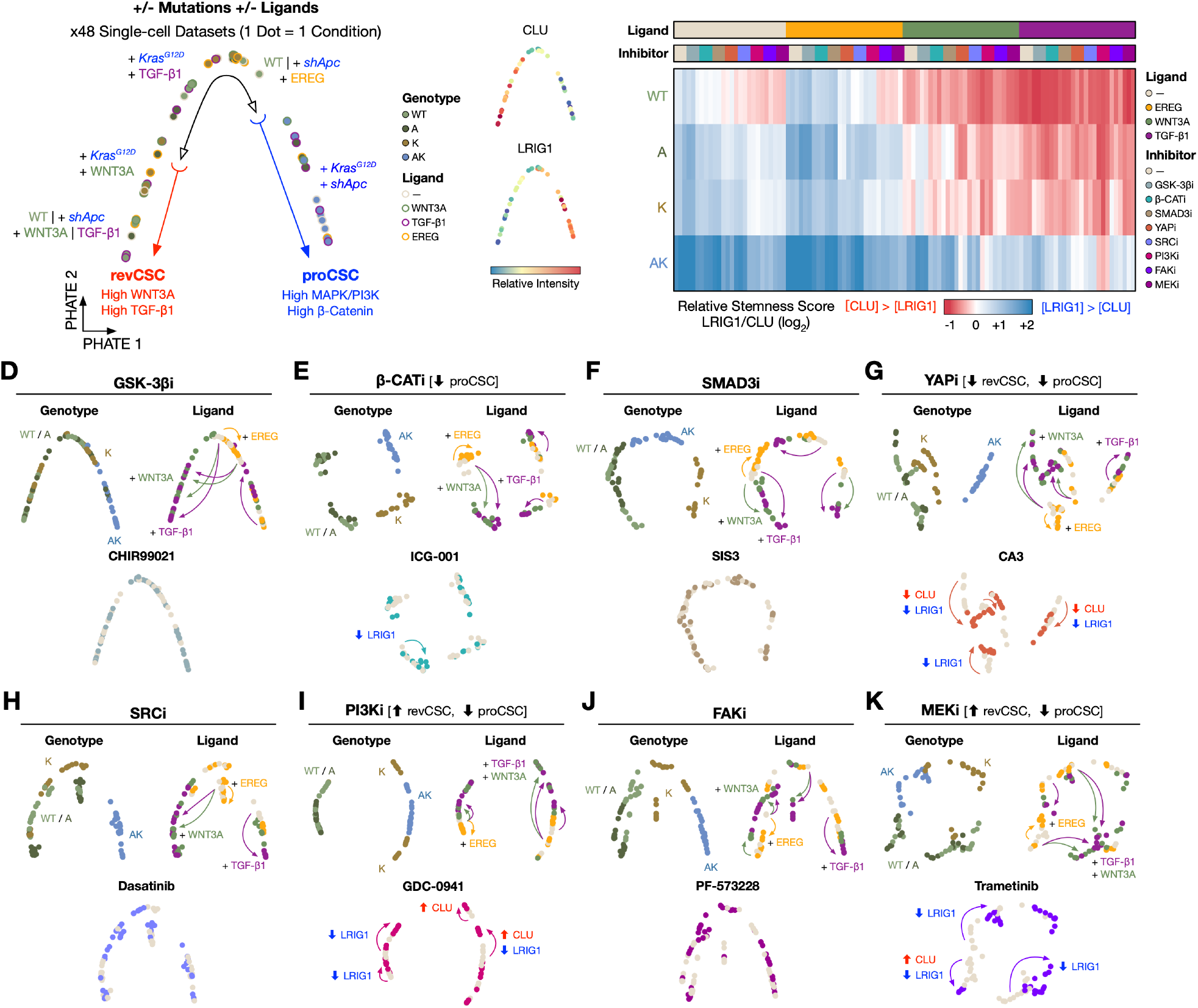
Signal Perturbation Analysis. Related to Figure 5. **A)** EMD-PHATE embedding of organoid cultures treated with ligands alone from the cue-signal-response array annotated with experimental metadata. One dot = one condition. **B)** PHATE embedding from **A)** coloured by EMD scores for CLU and LRIG1. **C)** Heatmap of relative stemness scores (log_2_-transformed single-cell expression ratio between LRIG1 and CLU) of 432 organoid cultures from the cue-signal-response array. **D-K)** EMDPHATE embeddings of organoid culture subsets from the cue-signal-response array focusing on each inhibitor.

**Figure S7.**
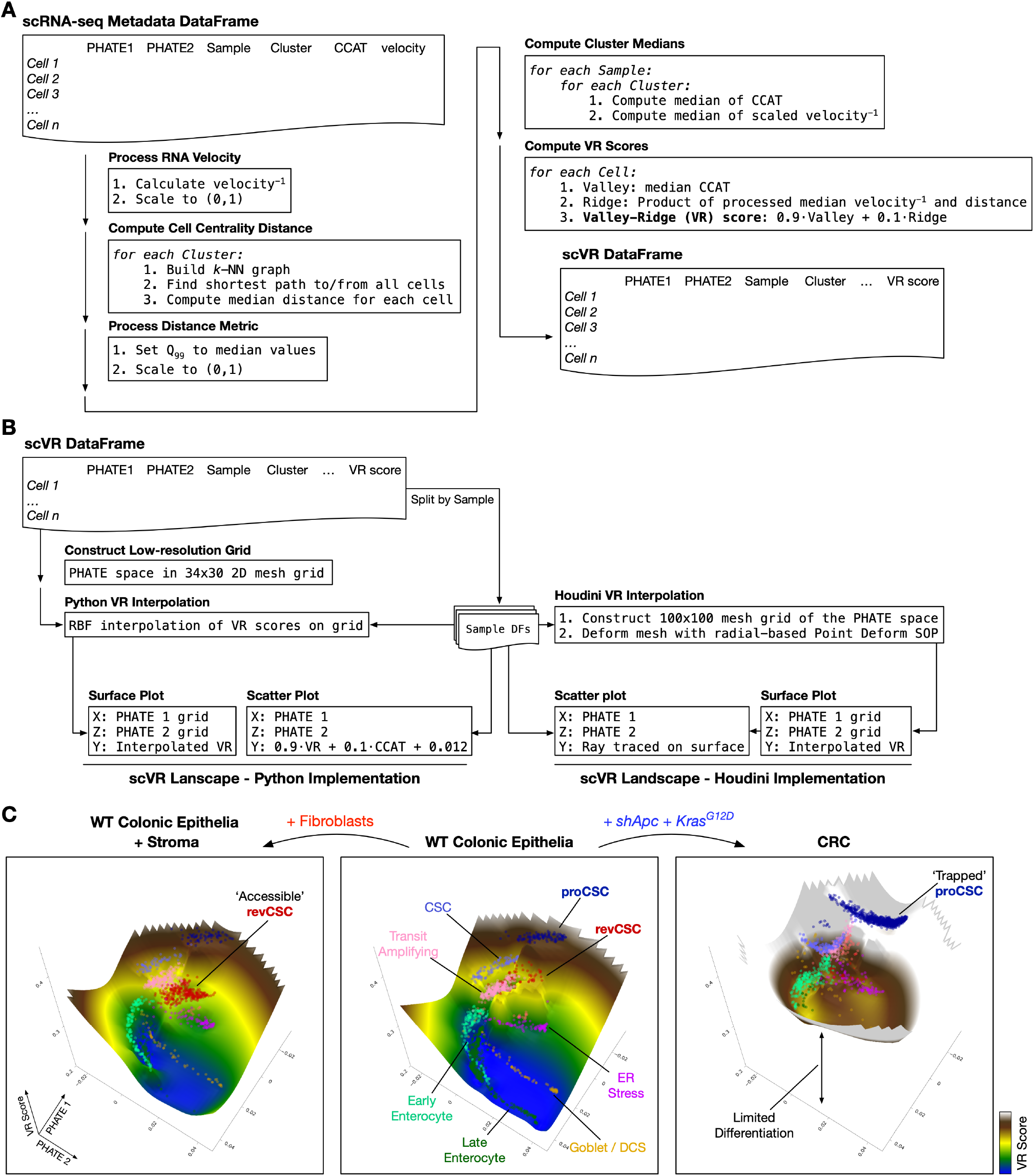
Generation of Waddington-like Landscapes from scRNA-seq Data. Related to Figure 6. **A)** Workflow for calculating VR scores from scRNA-seq data. **B)** Workflow for visualising PHATE and VR scores as 3D landscapes with either Python (left) or Houdini (right). RBF, radial basis function. **C)** PHATE and VR score landscapes visualised in Python.

## Notes

https://doi.org/10.5281/zenodo.7586958

https://community.cytobank.org/cytobank/experiments#project-id=1460

https://github.com/TAPE-Lab/Qin-CardosoRodriguez-et-al

https://www.ncbi.nlm.nih.gov/bioproject/PRJNA883610

### Competing Interest Statement

The authors have declared no competing interest.

