## Supplementary Table 1 for "A Single-cell Perturbation Landscape of Colonic Stem Cell Polarisation"

**Table S1. Organoid / Fibroblast Co-culture Mass Cytometry Antibody Panel**

| **Isotope-Metal** | **Antigen / Target** | **Antibody Clone** | **Supplier** |
| --- | --- | --- | --- |
| 89-Y | Phospho-Histone H3 [S28] | HTA28 | BioLegend |
| 106-Cd | RFP | 8E5.G7 | Rockland |
| 110-Cd | mCherry | 16D7 | Thermo |
| 111-Cd | CD44 | IM7 | BioLegend |
| 113-In | CD66a (CEACAM1) | CC1 | Thermo |
| 115-In | Pan-Cytokeratin (Pan-CK) | AE1/AE3 | BioLegend |
| 116-Cd | GFP | 5F12.4 | eBiosciences |
| 124-Te | TOB*is* Barcode | - | Prof. Mark Nitz |
| 126-Te | TOB*is* Barcode | - | Prof. Mark Nitz |
| 127-I | IdU (S-phase) | - | Fluidigm |
| 128-Te | TOB*is* Barcode | - | Prof. Mark Nitz |
| 130-Te | TOB*is* Barcode | - | Prof. Mark Nitz |
| 141-Pr | Phospho-PDPK1 [S241] | J66-653.44.22 | BD Biosciences |
| 142-Nd | Cleaved-Caspase 3 [D175] | D3E9 | CST |
| 143-Nd | C-MYC | D84C12 | CST |
| 144-Nd | Podoplanin (PDPN) | 8.1.1 | BioLegend |
| 145-Nd | Phospho-NDRG1 [T346] | D98G11 | CST |
| 146-Nd | Phospho-MKK4/SEK1 [S257] | C36C11 | CST |
| 147-Sm | Phospho-BTK [Y551] | 24a/BTK | BD Biosciences |
| 148-Nd | Phospho-SRC [Y418] | SC1T2M3 | BD Biosciences |
| 149-Sm | Phospho-4E-BP1 [T37/46] | 236B4 | CST |
| 150-Nd | Phospho-RB [S807/811] | J112-906 | BD Biosciences |
| 151-Eu | Phospho-PKCα [T497] | K14-984 | BD Biosciences |
| 152-Sm | Phospho-AKT [T308] | J1-223.371 | BD Biosciences |
| 153-Eu | Phospho-CREB [S133] | 87G3 | CST |
| 154-Sm | Phospho-SMAD1 [S463/465]  Phospho-SMAD5 [S463/465]  Phospho-SMAD9 [S465/467] | D5B10 | CST |
| 155-Gd | Phospho-AKT [S473] | D9E | CST |
| 156-Gd | Phospho-NF-κB p65 [S529] | K10-895.12.50 | BD Biosciences |
| 157-Gd | Phospho-MKK3 [S189] / MKK6 [S207] | D8E9 | CST |
| 158-Gd | Phospho-p38 MAPK [T180/Y182] | D3F9 | CST |
| 159-Tb | Phospho-MAPKAPK2 [T334] | 27B7 | CST |
| 160-Gd | Phospho-AMPKα [T172] | 40H9 | CST |
| 161-Dy | Phospho-BAD [S112] | 40A9 | CST |
| 162-Dy | LRIG1 | Polyclonal | R&D Systems |
| 163-Dy | Phospho-p90RSK [T359] | D1E9 | CST |
| 164-Dy | Phospho- p120-Catenin [T310] | 22/p120 (pT310) | BD Biosciences |
| 165-Ho | β-Catenin [Active] | D13A1 | CST |
| 166-Er | Phospho-GSK-3β [S9] | D85E12 | CST |
| 167-Er | Phospho-ERK1/2 [T202/Y204] | 20A | BD Biosciences |
| 168-Er | Phospho-SMAD2 [S465/467]  Phospho-SMAD3 [S423/425] | D27F4 | CST |
| 169-Tm | EPHB2 | 2H9 | BD Biosciences |
| 170-Er | Phospho-MEK1/2 [S221] | 166F8 | CST |
| 171-Yb | SOX9 | 3C10 | Abcam |
| 172-Yb | Cleaved-Caspase 8 [D387] | D5B2 | CST |
| 173-Yb | Cyclin B1 | GNS-11 | BD Biosciences |
| 174-Yb | Clusterin | Polyclonal | bio-techne |
| 175-Lu | Survivin | Polyclonal | bio-techne |
| 176-Yb | FABP2/I-FABP | 323730 | bio-techne |
| 191-Ir | DNA | - | Fluidigm |
| 193-Ir | DNA | - | Fluidigm |
| 194-Pt | Cisplatin (Dead Cells) | - | Fluidigm |
| 196-Pt | TOB*is* Barcode | - | Fluidigm |
| 198-Pt | TOB*is* Barcode | - | Fluidigm |
| 209-Bi | DiMeHH3 [K4] | C64G9 | CST |

Extracellular / Intracellular / Non-Antibody Parameter
